# DNA-foraging bacteria in the seafloor

**DOI:** 10.1101/528695

**Authors:** Kenneth Wasmund, Claus Pelikan, Margarete Watzka, Andreas Richter, Amy Noel, Casey R.J. Hubert, Thomas Rattei, Thilo Hofmann, Craig W. Herbold, Alexander Loy

## Abstract

Extracellular DNA is a major macromolecule in global element cycles, and is a particularly crucial phosphorus as well as nitrogen and carbon source for microorganisms in the seafloor. Nevertheless, the identities, ecophysiology and genetic features of key DNA-foraging microorganisms in marine sediments are completely unknown. Here we combined microcosm experiments, stable isotope probing and genome-centric metagenomics to study microbial catabolism of DNA and its sub-components in anoxic marine sediments. ^13^C-DNA added to sediment microcosms was degraded within ten days and mineralised to ^13^CO_2_. Stable isotope probing showed that diverse *Candidatus* Izemoplasma, *Lutibacter, Shewanella, Fusibacteraceae* and *Nitrincolaceae* incorporated DNA-derived ^13^C-carbon. Genomes representative of the ^13^C-labelled taxa were recovered and all encoded enzymatic repertoires for catabolism of DNA. Comparative genomics indicated that DNA can be digested by diverse members of the order *Candidatus* Izemoplasmatales (former *Tenericutes*), which appear to be specialised DNA-degraders that encode multiple extracellular nucleases. *Fusibacteraceae* lacked genes for extracellular nucleases but utilised various individual purine- and pyrimidine-based molecules, suggesting they ‘cheated’ on liberated sub-components of DNA. Close relatives of the DNA-degrading taxa are globally distributed in marine sediments, suggesting that these poorly understood taxa contribute widely to the key ecosystem function of degrading and recycling DNA in the seabed.

## Introduction

Subsurface environments including marine sediments harbour the bulk of microbial biomass on Earth^1^. Nevertheless, the functional capabilities and ecological roles of most of the diversity of subsurface microorganisms remain completely unknown^2^. The vast majority of microorganisms inhabiting marine sediments are ultimately sustained by heterotrophic metabolisms. Heterotrophs are predominantly fuelled by organic matter derived from primary production in the overlying water column and/or from land-derived inputs^3, 4^. Heterotrophic metabolisms are sustained by direct involvement in the initial break-down and catabolism of organic molecules, or indirectly, via the use of by-products of such catabolism, e.g., low molecular weight degradation products or fermentation products^5^. Additionally, cell debris released after the death and lysis of organisms, as well as excreted exometabolites, supply organic molecules for the growth of microorganisms^6, 7^. These sources collectively provide an immense diversity of organic molecules^8, 9^. Many of these organics can be used as nutrient and energy sources to sustain expansive numbers of microorganisms of very diverse composition in marine sediments^10^. The identities of microorganisms that use different kinds of organic molecules in marine sediments, are however, largely unknown. This is despite the fact that such knowledge lies central to our fundamental understandings of how microorganisms control biogeochemical cycles. It is also important for our understandings of ecological niche differentiation among microorganisms that utilise different substrates, and how this ultimately influences the diversity and community structures of microorganisms in the seafloor.

In general, the most abundant of the various classes of organic molecules available for catabolism by microorganisms in marine sediments, and in most other environments, include proteins, lipids, carbohydrates and nucleic acids – the molecules that make-up most of the biomass of any cell or virus^11-13^. Of these, DNA is known to contribute substantially to oceanic and sedimentary biogeochemical cycles, acting as a significant source of carbon, nitrogen and phosphorus, and providing an energy source^14-16^. Estimates suggest ‘bioavailable’ DNA, i.e., DNA available for digestion by extracellular nucleases, supplies microbial communities of coastal and deep-sea sediments with 2-4% of their carbon requirements, 7-4% of their nitrogen requirements, and a remarkable 20-47% of their phosphorus requirements^14, 16^. Analyses of microbial 16S rRNA genes derived from extracellular- or intracellular-DNA pools in marine sediments showed that extracellular DNA is rapidly turned-over in both shallow and deeper sediments (down to 10-12 meters)^17, 18^.

Some microorganisms are capable of catabolizing DNA via concerted steps involving extracellular digestion^19^, import systems^20^, and catabolic breakdown of imported sub-components in the cytoplasm^21^. Degradation products of nucleic acids such as urea and ammonium, as well as CO_2_ and acetate^22^, are also important nutrients for other members of microbial communities. Nucleic acids might be especially important phosphorus sources in sediments rich in metal-oxides such as Fe(III)- or Mn(IV)-oxides, whereby strong sorption of phosphorus to the metal oxides can occur, thereby diminishing bioavailable pools of this crucial nutrient^23^. Despite the fact that nutrient-rich and ubiquitous DNA biomolecules are available in marine sediments, our knowledge of microorganisms that degrade and mineralise DNA remains poor^24^.

In this study, we identified key-players that degrade and catabolise extracellular DNA in marine sediments. We hypothesized this abundant organic macromolecule might nourish and explain the presence of some of the commonly found taxonomic groups of microorganisms in marine sediments. To identify and study the ecophysiology of DNA-degrading microorganisms, we set-up microcosms with sediment from the Baffin Bay in between Canada and Greenland. Microcosms were either supplemented with ^13^C-labelled DNA, or individual DNA sub-components, and incubated under cold, anoxic conditions. We then analysed biogeochemical parameters and the microbial communities over a time-series of weeks via DNA-based stable isotope probing (DNA-SIP) and 16S rRNA gene amplicon sequencing. We subsequently retrieved the genomes of DNA-degraders identified by SIP via metagenomics to examine them for encoded DNA-degrading enzymes and catabolic pathways for molecular sub-components of DNA. The combined results provided multiple lines of evidence for the ability to degrade DNA by diverse groups of poorly understood bacteria, thereby providing new insights into potential niche occupation and *in situ* functions of these bacteria in the seafloor.

## Results

### Microcosm experiments show mineralization of DNA under cold, anoxic conditions

Anoxic microcosms with slurries of marine sediments from the Baffin Bay, Greenland, were individually amended with ^13^C-labelled or unlabelled genomic DNA from the archaeon *Halobacterium salinarum*, as well as the unlabelled nucleobases adenine, thymine, guanine and cytosine, and the nucleosides 2-deoxyadenosine and thymidine (Supp. Table 1). Complete mineralization of added DNA to CO_2_ was confirmed by isotopic analysis of CO_2_ in the headspace of the microcosms. This showed CO_2_ became enriched in ^13^C already at early time points (4 days), and that ^13^C-CO_2_ was produced over time (Supp. Fig. 1). Bacterial community analyses via amplicon sequencing of 16S rRNA genes from the microcosms revealed that the supplemented *Halobacterium* DNA was mostly depleted from approximately 1.3% to 0.1% from day 4 to day 10, respectively, and was only present in minor relative abundances (<0.1%) from day 13 and after (Supp. Fig. 2). Chemical analyses showed sulfate was not depleted at all and sulfide did not accumulate (results not shown), while high concentrations of total manganese and iron were measured that may support metal-reducing populations (Supporting information). Bacterial taxa related to known metal-reducing microorganisms^25^ collectively dominated both starting sediments and sediments in microcosms during the incubations. Bacterial community compositions among microcosms treated with DNA and controls were similar over the time of the experiment (Supporting information, Supp. Fig. 3 and 4).

**Fig. 1.**
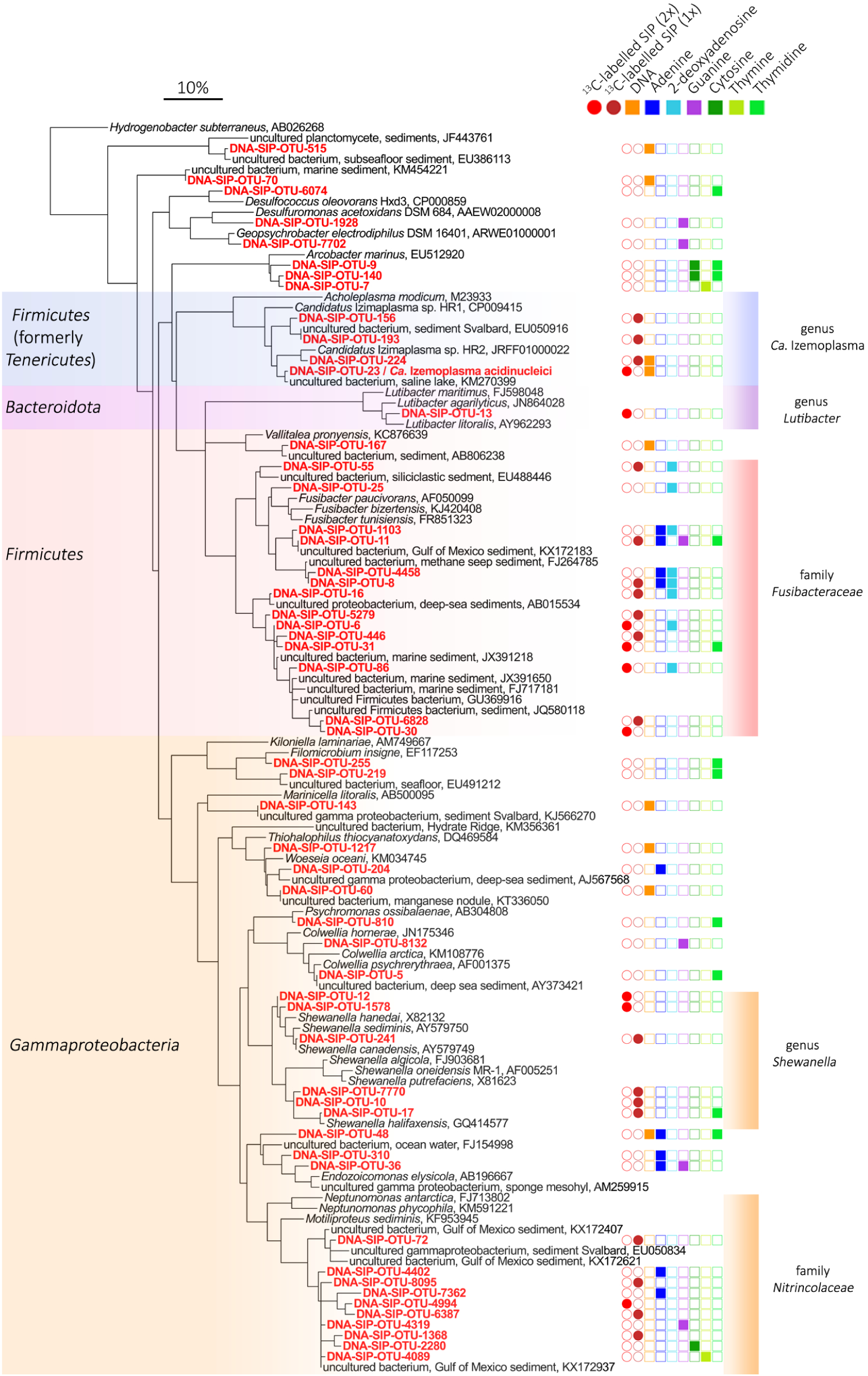
Phylogeny of 16S rRNA OTUs determined to be enriched in ^13^C in microcosms with ^13^C-DNA as substrate and/or to have significantly increased in relative gene abundance in microcosms with unlabelled DNA or selected DNA sub-components. OTUs that were labelled at two points (2x) are indicated with bright red, while relatives that were labelled at one time point (1x) are indicated in dark red. 16S rRNA OTU sequences recovered in this study are highlighted in red and bold. Taxonomic groups that included DNA-degrading bacteria as determined by DNA-SIP analyses are highlighted by shaded colours. Genbank accessions are included for 16S rRNA gene reference sequences. The scale bar represents 10% sequence divergence.

**Fig. 2.**
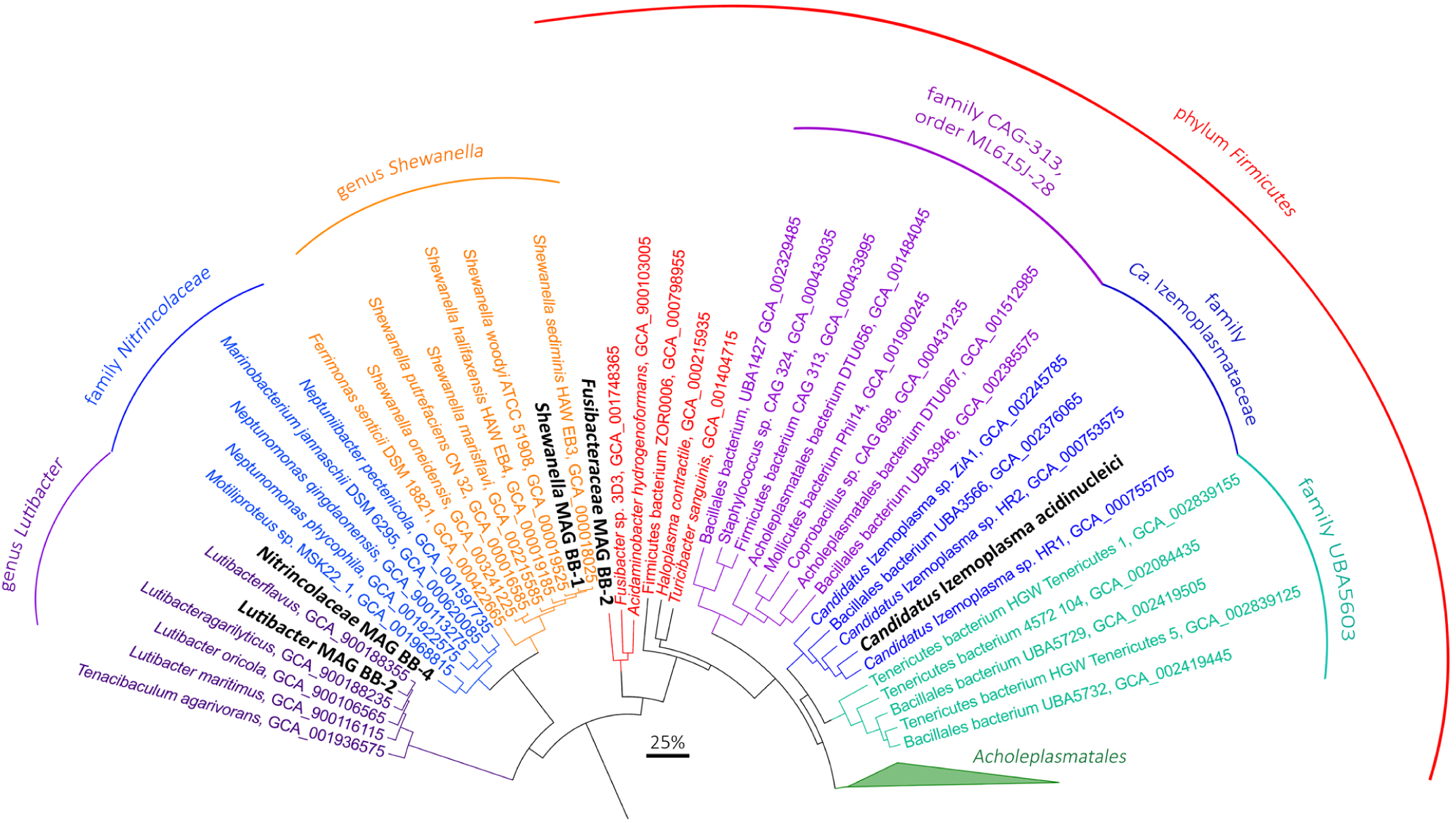
Genome-based phylogeny using concatenated protein sequences derived from single copy marker genes retrieved from CheckM analyses. MAGs recovered in this study are highlighted in black and bold. The approximate maximum-likelihood tree was calculated using FastTree. The scale bar represents 25% sequence divergence.

**Fig. 3.**
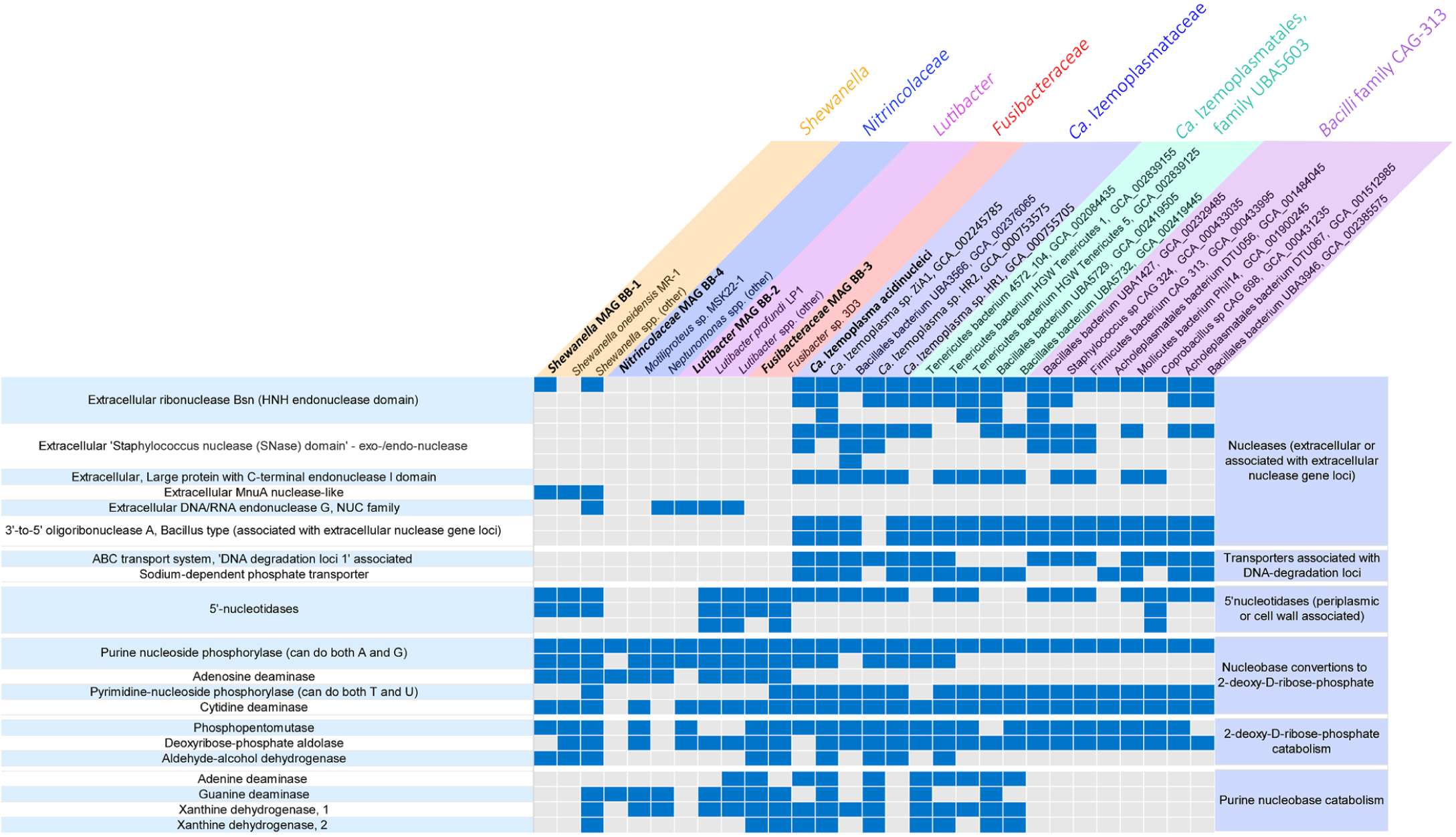
Presence-absence of key genes for enzymes involved in DNA degradation, transport and catabolism of DNA sub-components in MAGs and genomes of selected isolates. A list of key annotations from MAGs identified in this study are listed in Supp. Table 3.

**Fig. 4.**
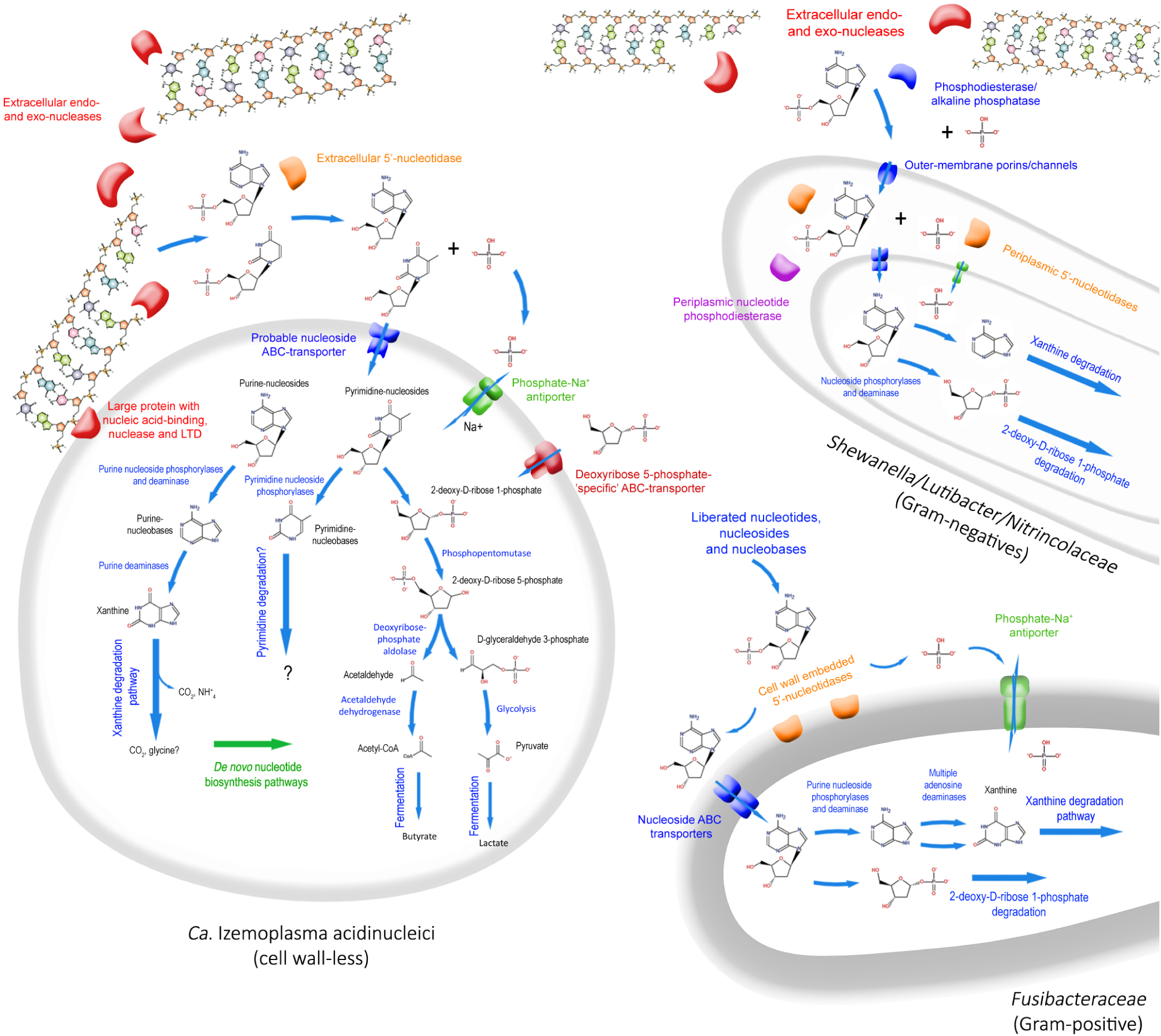
Taxon-centric schematic depiction of key enzymes and other functional proteins related to DNA catabolism and transport annotated from MAGs and close relatives. More detailed pathways for catabolism of DNA sub-components are provided for *Ca*. Izemoplasma acidinucleici, and for clarity, simplified versions of these redundant pathways are provided as arrows for the other organisms. LTD = Lamin tail domain. DNA molecule structures were sourced and modified from Wikipedia, and are under the Creative Commons license.

### SIP identifies diverse taxa that incorporate ^13^C-carbon from DNA

DNA-SIP identified bacterial taxa that incorporated ^13^C-carbon from the added ^13^C-DNA during replication and growth, and therefore must have either: i) catabolised the added DNA and/or sub-components of DNA, ii) salvaged nucleobases or nucleosides for incorporation during replication, or iii) utilised fermentation products released from catabolism of the added DNA. Numbers of bacterial 16S rRNA genes determined from qPCR analyses across all gradient density fractions revealed only slightly higher abundances of 16S rRNA genes in some ‘heavy’ density fractions (≥1.725 g ml^-1^)^26^ from incubations with ^13^C-DNA compared to ^12^C-DNA, from several time points, i.e., days 4, 10 and 13 (Supp. Fig. 5). Later time points showed no elevated 16S rRNA gene abundances in heavy fractions and therefore subsequent analyses focused on the first three time points, and thus, largely on the primary DNA degraders.

**Fig 5.**
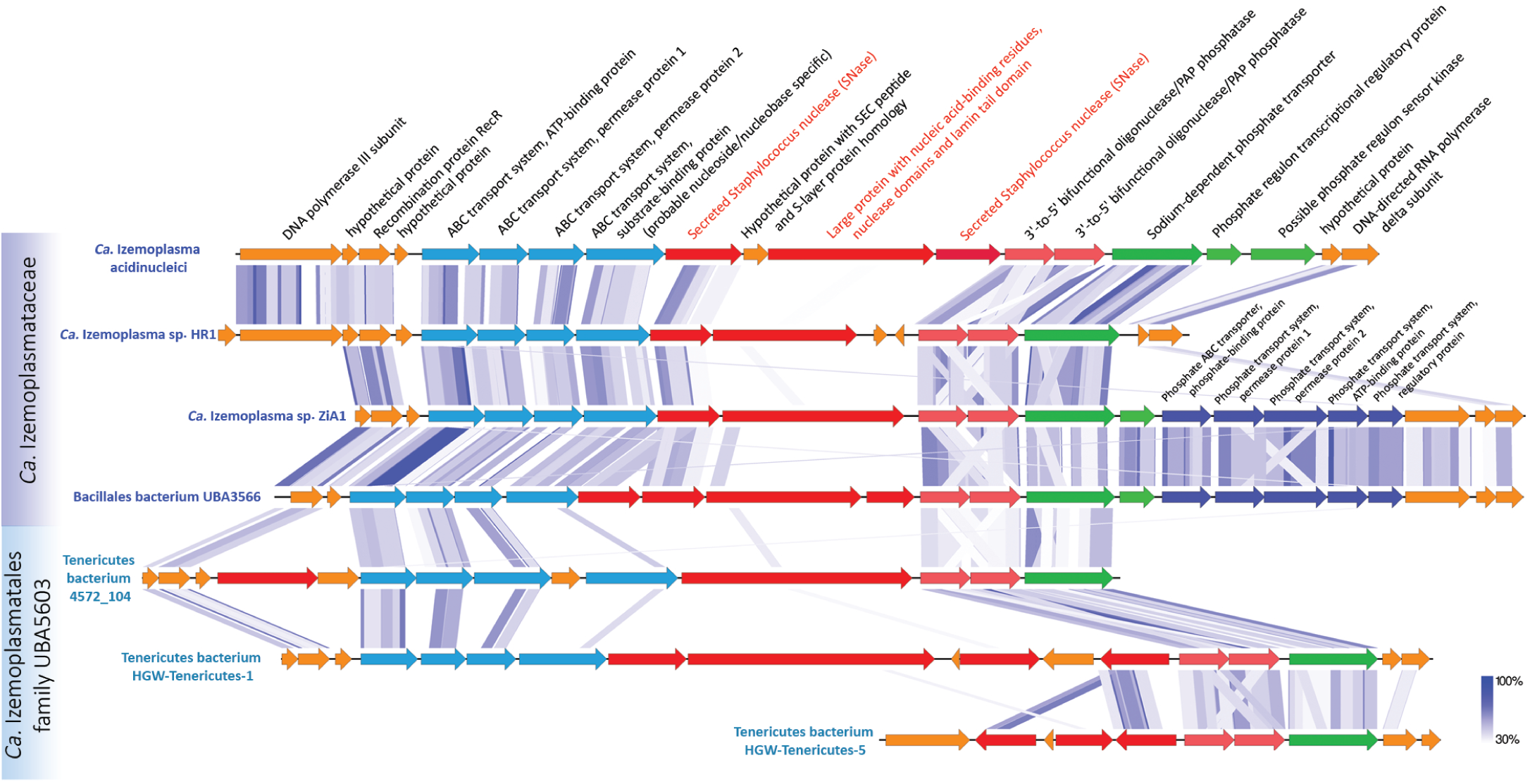
Gene arrangements of a putative ‘DNA-degradation loci’ in *Ca*. Izemoplasmatales MAGs. Shaded blue lines show regions with high sequence similarity and synteny as determined by tBLASTx using EasyFig. DNA-binding domains were predicted based on NCBI Conserved Domains (via BLASTP). Proteins predicted to be secreted are highlighted and labelled in red. Only scaffolds from MAGs with all or most genes present are included.

Amplicon sequencing of 16S rRNA genes across various CsCl_2_ density gradients identified individual operational taxonomic units (OTUs) that displayed significantly increased relative abundances in multiple ‘heavy’ density fractions (>1.725 g ml^-1^) from ^13^C-DNA treatments versus similar densities from ^12^C-DNA control treatments. We first identified individual OTUs that were significantly enriched at two or more time points. Further, we identified OTUs that were significantly enriched at only one time point but belonged to the same bacterial groups as those enriched at two time points. This identified 27 OTUs that belonged to five bacterial taxa (Fig. 1). The summed relative abundances of all OTUs from these groups were calculated across the different density gradients and showed significantly higher relative abundances of these taxa in ‘heavy’ fractions of the gradients (Supp. Fig. 6). OTUs related to the *Nitrincolaceae* did not collectively show higher relative abundances in heavy fractions from ^13^C-DNA treatments, except for OTU 4994 (Supp. Fig. 6). Phylogenetic analyses of 16S rRNA genes from the labelled OTUs showed that they were affiliated with: i) the genus *Shewanella* (class *Gammaproteobacteria*); ii) the family *Fusibacteraceae* (phylum *Firmicutes*), iii) the genus *Candidatus* Izemoplasma (GTDB phylum *Firmicutes*, formerly phylum *Tenericutes*); iv) the genus *Lutibacter* (GTDB phylum *Bacteroidota*, formerly phylum *Bacteroidetes*); and v) the family *Nitrincolaceae* (GTDB, formerly *Oceanospirillaceae*) (class *Gammaproteobacteria*) (Fig. 1). The naming of the *Ca.* Izemoplasma (formely *Ca*. Izimaplasma) is described in further detail below. Examination of 16S rRNA gene sequences in publically available sequence databases showed that closely related sequences from these five groups are globally distributed in coastal and deep-sea sediments (Supporting information, Supp. Fig. 7).

### Distinct responses of diverse bacteria to additions of DNA, nucleobases or nucleosides

Complementary to the SIP analyses, we also tracked the relative abundances of individual OTUs in response to additions of DNA, as well as additions of individual sub-components of DNA (Fig. 1 and Supp. Fig. 8), i.e., the nucleobases adenine, thymine, guanine and cytosine, and the nucleosides 2-deoxyadenosine and thymidine (Supp. Table 1). This was performed in order to obtain indications for growth in response to the addition of these substrates. Among the OTUs that were identified as incorporating ^13^C-carbon from ^13^C-DNA by SIP analyses, two *Ca.* Izemoplasmataceae OTUs (23 and 224) showed significant increases in relative abundances in microcosms with DNA additions versus no-substrate controls, over multiple time points (Supp. Fig. 8). For nucleobases, only 5 OTUs responded to the pyrimidines thymine or cytosine, while 14 OTUs responded to the purines adenine or guanine (Fig. 1). Compared to thymine, to which only 2 OTUs responded, 11 different OTUs responded to thymidine. Interestingly, although no *Fusibacteraceae* OTUs became enriched in microcosms in response to additions of DNA only, numerous OTUs (n=9) from this group were enriched in response to purine-based nucleobases or nucleosides, especially those containing adenine moieties. Six of these *Fusibacteraceae* OTUs were determined to be ^13^C-labelled in the SIP experiment. *Fusibacteraceae* OTUs were the only taxa enriched by 2-deoxyadenosine, whereby 8 different OTUs responded. For instance, *Fusibacteraceae* OTU 8 dominated communities amended with 2-deoxyadenosine at day 10, reaching approximately 37% in relative abundance, versus 1% in the no-substrate controls (Supp. Fig. 8). Four of the *Fusibacteraceae* OTUs that responded to 2-deoxyadenosine also responded to adenine. These four OTUs also showed higher responses to the sugar-containing nucleoside 2-deoxyadenosine than the corresponding nucleobase adenine. Apart from taxa that were also labelled in the SIP analyses, several OTUs of the family *Endozoicomonadaceae* (class *Gammaproteobacteria*) were noticeably enriched in a number of treatments in this study, i.e., by DNA, adenine, guanine and thymidine (Fig. 1 and Supp. Fig. 8).

### Recovery of genomes of DNA-degrading taxa

Near complete (>90%) metagenome-assembled genomes (MAG) that were representative of the taxa labelled in the SIP experiments were recovered (Supp. Table 2). These MAGs together with other publically available genomes and MAGs from related bacteria^23, 27-31^, were analysed by phylogenomics (Fig. 2) and for genes encoding enzymes required to catabolise DNA (Fig. 3 and Supp. Table 3). The *Ca.* Izemoplasma MAG contained a 16S rRNA gene sequence that was 100% identical to OTU 23, which was labelled in the SIP experiment. All other MAGs recovered here did not contain 16S rRNA gene sequences. The *Nitrincolaceae* MAG BB-4 was predicted to be 79% complete. This MAG may not be representative of the *Nitrincolaceae* OTU 4994 that was determined to be labelled in the SIP analyses, since that OTU was present in very low abundance (<0.03% in any microcosm) and was unlikely to have been assembled and binned by our sequencing effort. Phylogenomic analyses (Fig. 2) and average nucleotide identity (ANI) analyses (Supp. Table 2), revealed that all MAGs constituted novel species or genera within their respective phylogenetic groups. For the *Ca.* Izemoplasma MAG, we propose the species name *Ca.* Izemoplasma acidinucleici (see below for detailed descriptions and Supporting information).

### Extracellular and cell wall associated nucleases encoded in DNA-degraders

A key determinant for the ability to utilise DNA as a nutrient and/or energy source is the ability for extracellular digestion of DNA polymers. Importantly, extracellular nucleases can also be differentiated from nucleases used for recycling nucleic acids in the cytoplasm^19^. Accordingly, we identified genes for extracellular nucleases in MAGs and/or related reference genomes of *Shewanella, Lutibacter, Nitrincolaceae* and *Ca.* Izemoplasma populations (Fig. 3 and Fig. 4). We did not, however, find any genes related to extracellular nucleases in the *Fusibacteraceae* MAG BB-3 or reference genomes. Strikingly, up to five extracellular nucleases were encoded in some of the marine *Ca.* Izemoplasmataceae (Fig. 3). Further, related MAGs from an undescribed *Ca.* Izemoplasmatales family UBA5603 (Fig. 2) and a related yet undescribed *Bacilli* family CAG-313 (Fig. 2), also harboured numerous copies of extracellular nuclease genes (Fig. 3). Encoded extracellular nucleases identified in our *Shewanella* MAG BB-2 are homologs to extracellular nucleases previously shown to be highly expressed when *Shewanella oneidensis* MR-1 was grown on DNA^23, 32^.

To facilitate the import of DNA sub-components into cells, further deconstruction of nucleotides and nucleosides is necessary^5^. Interestingly, MAGs from the *Ca.* Izemoplasmataceae and *Ca*. Izemoplasmatales family UBA5603 harboured conspicuous genomic loci associated with multiple extracellular nuclease genes that encoded enzymes that may facilitate further digestion and import of nucleic acid components, i.e., additional 3’-5’ nucleases, phosphohydrolases, ABC-transporters and phosphate transporters (Fig. 5). Sequence homology searches of the ‘substrate-binding subunit’ of the ABC-transporters against the IMG database^33^ identified that the genomes of organisms with best hits also encoded nucleases or nucleosidases within the same genomic neighbourhood. This indicated that these are probable nucleoside or nucleobase transporters. Additionally, this loci encoded a large protein (1049 amino acids) (Fig. 5) that was noteworthy because it contained nucleic acid-binding, nuclease and ‘lamin-tail’ domains (Supp. Fig. 11 and see Supp. Results and Discussion).

The *Shewanella* MAG BB-2 recovered here encoded several nucleotidases previously shown to be critical for growth on extracellular DNA^23^, i.e., excreted bifunctional 2’,3’ cyclic nucleotide 2’ phosphodiesterase/3’ nucleotidases and a putative UDP-sugar hydrolase protein with 5’ nucleotidase domains (included under nucleotidases in Fig. 3). *Lutibacter* and *Shewanella* also harboured multiple copies of genes for 5’-nucleotidases that were predicted to the reside in their periplasms (Fig. 3 and Fig. 4). Similarly, *Fusibacteraceae* and *Ca*. Izemoplasmataceae genomes encoded 5’-nucleotidases that were predicted to be located in their cell walls or extracellularly, respectively (Fig. 3 and Fig. 4). Dephosphorylation of nucleotides by nucleotidases is critical because cell membranes are impermeable to nucleotides due to negatively-charged phosphate groups^34, 35^.

### Catabolic pathways for sub-components of DNA

Once imported into the cytoplasm, purine- and pyrimidine-deoxyribonucleosides may be further broken-down, whereby the respective bases may then enter catabolic pathways or salvage pathways used for incorporation into newly synthesised nucleic acids. We identified two copies of predicted ‘purine deoxyribonucleoside phosphorylases’ in most genomes of putative DNA-degraders (Fig. 3). The two different copies presumably offer specificity to the two purine nucleobases adenine and guanine. Specific adenosine deaminases were also annotated in all genomes of putative DNA-degraders, except for any of the *Ca*. Izemoplasmataceae. The *Fusibacter* sp. 3D3 reference genome harbored multiple copies of adenosine deaminase genes. Genes for ‘pyrimidine deoxyribonucleoside phosphorylases’ were detected in most *Ca*. Izemoplasmataceae genomes and in the *Fusibacter* sp. 3D3 reference genome. All putative DNA-degrading taxa encoded cytidine deamidases. Although uracil-specific phosphorylases that are required for further processing of 2-deoxycytidine via 2-deoxyuracil could not be identified, it is possible that the pyrimidine-nucleoside phosphorylases can catalyse both due to their known bifunctionality for these structurally similar substrates^36^. All DNA-degraders identified in this study therefore have enzymes required to cleave nucleobases into the sugar and base components.

The potential to catabolise the purine bases adenine and guanine were indicated by several lines of evidence. Genes for deaminases specific for both purine bases were present in *Fusibacteraceae* and various *Ca*. Izemoplasmataceae genomes (Fig. 3). In *Lutibacter* spp., adenine deaminase genes could only be identified in two organisms in the NCBI database, while in *Shewanella* spp., only *S. halifaxensis* appeared to harbor this gene. Guanine deaminase genes were also present in numerous *Lutibacter* and *Nitrincolaceae*, and were only identified in one *Shewanella*, i.e., *S*. *algae*.

In MAGs of the marine *Ca*. Izemoplasmataceae and the sister family UBA5603 (order *Ca.* Izemoplasmatales), genes for the two purine-specific deaminases were often located in close vicinity to each other, and were distinctively located in the same genomic region as genes for xanthine dehydrogenases and their accessory proteins (Supp. Fig. 9). Xanthine dehydrogenases are key enzymes required for conversions of purine bases to xanthine, which is a common intermediate of purine breakdown^37^. These genomic loci were highly similar in *Ca*. Izemoplasmatales and the genomes of some *Clostridium* species (Supp. Fig. 9), which are among the few biochemically characterised anaerobic purine-degraders^21, 37^. Genes for enzymes required for the further catabolism of xanthine could not be unambiguously identified, since most are not described and are only predicted to occur based on biochemical inferences^21^. Nevertheless, several enzymes predicted to have amidohydrolase and hydantoinase activities were also encoded between the xanthine dehydrogenase genes (Supp. Fig. 9). These possibly play a role in the cleavage of the heterocyclic rings, and thus, further catabolism of xanthine. Genes for any enzymes involved in purine catabolism were notably absent from any MAGs from the *Bacilli* family CAG-313 (Fig. 3).

Apart from indications for the dephosphorylation of pyrimidines described above, we were unable to identify genes for anaerobic catabolic degradation of pyrimidine bases. This was not possible because the enzymes involved in this process in anaerobic microorganisms have not been identified.

Importantly, all MAGs representative of taxa that became labelled in our SIP experiment had the potential to catabolise the 2-deoxy-D-ribose-phosphate sugar moiety that can be liberated from nucleosides. In this regard, we identified genes for enzymes for catabolism of 2-deoxy-D-ribose-phosphate to D-glyceraldehyde 3-phosphate, an intermediate of glycolysis, and acetaldehyde, in all our MAGs (Fig. 3 and Fig. 4). Genes for aldehyde-alcohol dehydrogenases that could convert acetaldehyde to acetyl-CoA, were present in most labelled groups except in *Lutibacter* and *Nitrincolaceae*, and were only present in some marine *Ca*. Izemoplasmataceae MAGs, i.e., *Ca*. Izemoplasma sp. ZiA1 and HR1. Genes for a protein microcompartment that was postulated to be used for catabolising the liberated aldehydes in *Ca*. Izemoplasma sp. HR2^27^, were not found in any other *Ca*. Izemoplasma MAGs analysed here.

### De novo biosynthetic pathways for nucleotides encoded in DNA-degrading taxa

We identified large complements of genes encoding enzymes involved in the complete *de novo* syntheses of nucleotides in the genomes of most *Ca*. Izemoplasma (Supp. Table 4) (see Supporting information for descriptions). All other DNA-degraders identified in this study belong to bacterial groups that do not require supplementation of nucleic acid building blocks to growth media^29, 38-40^.

### Proposal of the novel species *Ca.* Izemoplasma acidinucleici and amendment of the genus name *Ca.* Izimaplasma to *Ca.* Izemoplasma

Based on a combination of unique phylogeny (based on 16S rRNA genes and concatenated single copy marker proteins) and genomic similarity (ANI), as well as genome-based predictions of nucleic acid catabolism that are supported by SIP results, we propose a novel species name *Candidatus* Izemoplasma acidinucleici for the *Ca*. Izemoplasma MAG recovered in this study. We have revised the previously suggested genus name *Ca*. Izimaplasma^27^ to *Candidatus* Izemoplasma (Bernhard Schink and Aharon Oren, personal communication) of the family *Izemoplasmataceae* and order *Izemoplasmatales*. We propose the MAG of *Candidatus* Izemoplasma acidinucleici as type material for this lineage, since it fulfils the properties outlined by Chuvochina *et*. *al*.^41^, with the only exception being that it is contained within 191 contigs.

- *Ca*. Izemoplasma gen. nov. (I.ze.mo.plas’ma. Gr. n. Izema, that what settles, sediment; - o-, connecting vowel; Gr. neut. n. plasma, something that forms, N.L. neut. n. Izemoplasma, a formed structure from sediment) with *Ca*. Izemoplasma acidinucleici (a.ci.di.nu.cle’i.ci. N.L. n. acidum nucleicu m, nucleic acid; N.L. gen. n. acidinucleici, of nucleic acid, referring to).

## Discussion

This study revealed the identities of key microbial players involved in the turn-over and recycling of extracellular DNA in anoxic marine sediments for the first time using culture-independent approaches. Since over 90% of the seafloor is permanently under 4°C^42^, the cold conditions used in our microcosm experiments represent conditions covering most of the seafloor. This was reflected by the detection of relatives of the DNA-degrading taxa in various sediments around the world’s oceans, suggesting that these DNA-degrading taxa may contribute to DNA turn-over and mineralization on a global scale. Such heterotrophic processes may also be especially important in future arctic systems, where increased primary production due to decreased ice-cover is expected to dramatically alter carbon cycles and possibly increase inputs of phyto-detrital organic matter to the seafloor^43^.

Members of the *Ca.* Izemoplasmataceae particularly stood out in this study because they became highly labelled in the SIP analyses, significantly increased in relative abundances when DNA was added to sediments and harboured various genes strongly indicating that they could catabolically exploit various sub-components of DNA. Strikingly, some *Ca.* Izemoplasmataceae encoded up to five extracellular nucleases, providing a clear indication that they can digest environmental DNA. Some of the extracellular nuclease genes were directly co-localised with genes encoding enzymes for further digestion of DNA into oligonucleotides and nucleosides, as well as removal of phosphates from nucleotides, and for transport of liberated nucleobases and phosphate into the cells. Together, the genomic co-localisations of these genes suggested that such processes are carried out in a co-ordinated manner in the cell and are therefore important nutrient acquisition strategies for these organisms.

Up to now, little was known regarding the ecological or biogeochemical roles of *Ca.* Izemoplasmataceae in marine sediments. One recent study of MAGs derived from marine sediments hinted that *Ca.* Izemoplasmataceae may catabolise nucleotides and the sugar moiety of DNA, among other key features such as the ability to grow via fermentation of few simple sugars^27^. Our analyses expand the ecophysiological understanding of *Ca.* Izemoplasmataceae by showing that they actively participate in the primary degradation of extracellular DNA polymers in anoxic marine sediments. The genomic analyses also show that they are evolutionary adapted for DNA catabolism, since they have relatively small genomes and limited catabolic repertoire for organics other than DNA. We therefore propose that they may establish their primary niche in sediments via the use of DNA as a nutrient and energy source. We also show that DNA-degrading capabilities are present in the *Ca*. Izemoplasmatales family UBA5603, which appears to be a sister clade of the marine *Ca.* Izemoplasmataceae that is primarily derived from the terrestrial subsurface. Genomes of the UBA5603 family (order Izemoplasmatales) did, however, lack many genes for enzymes required for *de novo* nucleotide biosynthesis, suggesting that they may degrade DNA for either, or both, salvage and/or catabolic purposes.

The detection of ^13^C-labelled *Shewanella* and *Lutibacter* in the SIP analyses served as a good indication that *bone fide* DNA-degrading bacteria were detected by our SIP approach. This is because *Shewanella* are known to grow by catabolising DNA as a sole nutrient and energy source *in vitro*^23^. Similarly, members of the genus *Lutibacter* have been shown to possess hydrolytic activity for DNA *in vitro*, although it has not been reported whether they can grow using DNA as a sole nutrient^39^. *Shewanella* and *Lutibacter* are known to be capable of catabolising various simple organic substrates *in vitro*, e.g., amino acids, carboxylates and sugars^28, 44^. Our data shows that diverse representatives of the *Shewanella* and *Lutibacter* also degrade DNA within *in situ*-like conditions. It further shows that they effectively compete with other DNA-degraders for DNA-derived carbon within the context of a complex community. This therefore provides important confirmation that these functional properties are expressed *in situ* and that they may be important nutrient acquisition strategies for these bacteria in marine sediments.

Our SIP analyses also showed various OTUs from the family *Nitrincolaceae* sourced carbon from DNA. We identified an extracellular nuclease encoded only in the genome of the isolate *Neptunomonas phycophila*, and accordingly, other enzymes required for catabolism of DNA-subcomponents were sparsely encoded among so far sequenced genomes of this family. Together, this suggested that degradation of extracellular DNA is a specialised physiological feature of only some *Nitrincolaceae* members. Indeed, various members of *Nitrincolaceae* are known to be a relatively versatile in catabolic terms, whereby they are known to degrade various carbohydrates, alcohols, organic acids and complex polycyclic aromatic hydrocarbons^45^. Alternatively, the low abundance *Nitrincolaceae* detected in this study and that could not be sampled for genomically, may have distinct genomes that harbour more expansive gene complements for enabling DNA catabolism.

Bacteria affiliated with the family *Fusibacteraceae* were conspicuous for their strong responses to additions of individual purine-based nucleobases and nucleosides, while also being highly labelled in the SIP analyses. Intriguingly, we could not identify known extracellular nucleases in the *Fusibacteraceae* MAG BB-3 or in any close relatives, and they did not increase in relative abundances when DNA was added as substrate. Although unknown extracellular nucleases may exist, their absence was notable because most previously described extracellular nucleases of anaerobic bacteria are derived from related *Firmicutes*, such as *Bacillus* and *Staphylococcus* species^46, 47^. This led us to infer that the *Fusibacteraceae* indeed lack extracellular nucleases. Instead, we could identify genes for multiple 5’-nucleotidases that were predicted to be embedded on their cell walls, as well as multiple gene clusters predicted to encode specific nucleoside/nucleotide transporters. Further, in-line with observations from the microcosm experiments that showed strong responses of multiple *Fusibacteraceae* OTUs to 2-deoxyadenosine amendments in the microcosms, the *Fusibacter* sp. 3D3 reference genome harboured multiple copies of adenosine deaminase genes. This may enable extra production of this enzyme and therefore more efficient processing of these molecules to the sugar and base moieties, which could then be further catabolised. Together, this indicates that these organisms may be ‘cheaters’, i.e., they may use sub-components of DNA liberated into the sedimentary environment by other DNA-degraders. *Fusibacter* isolates are generally known to be capable of utilizing various simple carbohydrates^48-50^, but until now, no *Fusibacter* species or closely related bacteria have been reported to have any activity related to DNA degradation or the degradation of molecular sub-components of DNA.

Finally, our analyses showed that additions of different individual sub-components of DNA to microcosms invoked highly distinct shifts in 16S rRNA gene relative abundances of diverse bacterial taxa. Interestingly, the different responses also occurred among distinct yet related members of some phylogenetic clades. For instance, various OTUs from the *Fusibacteraceae* responded differently to several DNA sub-components, especially purine-based molecules. This shows how different sub-components of DNA have the potential to be partitioned among different microorganisms. Free DNA subcomponents could arise if they are liberated from the parent molecule, e.g., after extracellular digestion of DNA by nucleases excreted by other primary DNA-degraders. This may therefore open-up multiple niches for different taxa, and in doing so, promote microbial diversity. In relation to this, additions of purine-based molecules consistently elucidated greater responses and from more diverse taxa, compared to responses to pyrimidine-based molecules. This aligns with previous research in estuarine waters, which showed that purine additions to water samples elicited considerably stronger end-product (urea) production than pyrimidines^51^. Further, we noted that more OTUs responded to nucleosides than their respective nucleobases. This signifies that the sugar moiety was preferentially exploited by some bacteria for energy conservation, rather than the base. This was especially notable for the pyrimidine nucleoside thymidine, which stimulated responses from eleven different OTUs, in comparison thymine itself that elucidated responses from only two OTUs.

## Conclusions

In this study, we provide complementary evidence supporting the involvement of diverse bacteria belonging to the *Lutibacter, Shewanella, Fusibacteraceae, Ca.* Izemoplasmataceae and *Nitrincolaceae* in the anaerobic degradation of DNA in marine sediments. The *Ca.* Izemoplasmataceae organisms are replete with genes for enzymatic machinery necessary to deconstruct DNA from the extracellular environment and to subsequently import and fully catabolise different molecular sub-components of DNA. These genes were often conspicuously co-localised in *Ca.* Izemoplasmataceae genomes, suggesting they have evolved to be regulated and function in a co-ordinated manner. Bacteria related to the *Fusibacteraceae* appeared to be very efficient at degrading purine-based DNA sub-components, but not extracellular DNA. We therefore propose that they may have been ‘DNA cheaters’ that utilised sub-components of DNA liberated by other members of the community. This study also supports results from previous *in vitro* studies of *Shewanella* and *Lutibacter* that demonstrated DNA-degrading capabilities, and importantly, shows that these bacteria perform such activities under *in situ*-like conditions. Together, this study sheds new light on the functional properties of several bacterial groups that are widespread yet poorly understood in marine sediments. These bacteria may thus collectively contribute to the essential process of turning-over the copious quantities of extracellular DNA that are present in vast expanses of the seafloor.

## Materials and Methods

### Sediment sampling and microcosm set-ups

Marine sediments (0-30 cm) were collected from two locations in northern Baffin Bay (Station ‘KANE-2B’ 79°30.909 −70°50.819 and ‘GL-111’ 76°18.354 −73°13.180) using a box corer aboard the Canadian Coast Guard icebreaker CCGS *Amundsen*, in July 2014. Sediments were filled into 250 ml Schott bottles to exclude oxygen, sealed and maintained at 4°C or on ice during transport. Sediments from both sites were mixed together in equal volumes, and then amended with cold (4°C) anoxic artificial seawater (1:1 v/v) (detailed in Supporting information). Resulting sediment slurries were and homogenised thoroughly within an anoxic glove-box (containing approximately 88% nitrogen, 2% hydrogen, 10% CO_2_). Approximately 30 ml of sediment slurry was added to 100 ml serum flasks, and these were sealed and crimped with 2cm thick black butyl rubber stoppers. Sediment slurries and microcosms were placed on top of frozen ice-packs during all preparation or manipulation steps in the anoxic glove-box. Microcosms were maintained at 4°C during all subsequent incubations.

### Microcosm incubation conditions and subsampling

Microcosms were pre-incubated for 4 days prior to substrate additions and this time point where substrates were added is herein referred to as day 0. Substrate were also added directly after subsampling at day 10. Purified ^13^C-labelled or unlabelled DNA extracted from *Halobacterium salinarum* (Supporting information), which is not present at detectable abundances in typical marine sediments, was added as substrate at concentrations of 36 µg ml^-1^. This amount was added because it is similar to that of total bioavailable DNA previously measured in marine sediments^14^. The nucleobases and nucleosides adenine, guanine, cytosine, thymine, 2-deoxyadenosine and thymidine (Sigma Aldrich, >99% pure) were added individually to separate microcosms, each to final concentration of 100 µM. Parallel microcosms without added substrates were analysed as ‘no-substrate’ controls. All treatment series (Supp. Table 1) were performed in triplicate microcosms.

Initial samples were taken directly after preparation of the sediment slurry, and following substrate amendment, subsamples were taken for microbial community profiling, SIP and metagenomics at days 0, 4, 10, 13, 24, and 31. For microbial community profiling via 16S rRNA gene amplicon sequencing, 100 µl from each microcosm was subsampled. Additional 1 ml subsamples were taken at each time point for chemical analyses. Subsamples were kept on ice-packs inside the anoxic glove-box during subsampling, and were then immediately snap-frozen on dry-ice outside the glove-box and subsequently stored at −80°C. All tubes for subsampling were introduced into the anoxic glove-box at least 1 hr prior to subsampling to remove traces of O_2_. Headspace gas (15 ml) was subsampled from microcosms using N_2_-flushed syringes fitted with needles and injected into pre-evacuated 12 ml glass exetainers (Labco, UK) that were sealed with butyl-rubber stoppers.

### DNA extractions

DNA for 16S rRNA gene amplicon sequencing and for density gradient centrifugations was extracted from 100 µl of microcosm slurries using a combination of bead beating, a cetyltrimethylammonium bromide-containing buffer and phenol-chloroform extractions (detailed in Supporting information). DNA for metagenomic library preparation was purified from 100 µl of microcosm subsamples using a MoBio Ultra Pure Power Soil Kit (MoBio, USA) following the manufacturer’s instructions.

### Density gradient centrifugations and quantitative PCR analyses of fractions

DNA from triplicates of specific treatments and time points to be analysed by DNA-SIP were combined equally to a total of 500 ng and subject to ultracentrifugation in a CsCl_2_ solution with a VTi-90 rotor for 72 hrs following standard protocols^52^. Gradients were collected by fractionation into approximately 20 separate fractions of 200 μl each. The densities of the collected fractions were measured using a refractometer (AR200; Reichert Analytical Instruments, USA) at 22°C. DNA was concentrated and purified by precipitation with polyethyleneglycol (MW=8000) and glycogen, and subsequently washed using methods previously described^52^. DNA was diluted 1:100 in ddH_2_O to facilitate PCR amplification.

### PCR and quantitative real-time PCR amplification of 16S rRNA genes

PCR amplifications of 16S rRNA genes from DNA extracted from microcosms and density gradient centrifugation fractions were performed using a two-step PCR barcoding approach^53^, using primers that target most *Bacteria* (341F: 5’-CCTACGGGNGGCWGCAG-3’ and 785R: 5’-GACTACHVGGGTATCTAATCC-3’)^54^. Both primers included ‘head’ sequences to facilitate barcoding in the second-step PCR^53^. PCR conditions are detailed in the Supporting information. PCR products were purified using a ZR-96 DNA Clean-up Kit(tm) (Zymo Research, USA) and pooled to equimolar concentrations after quantification using the Quant-iT(tm) PicoGreen® dsDNA Assay Kit (ThermoFischer Scientific, Germany). Sequencing was performed by MicroSynth (Switzerland) on an Illumina MiSeq instrument using MiSeq Reagent Kit v3 chemistry with 300 bp paired-end read mode. Negative DNA extract controls (no added sediment) and PCR negative controls (no DNA template) were performed and sequenced in parallel (even if no PCR product was detected).

Quantitative real-time PCR assays for bacterial 16S rRNA genes from fractions collected from CsCl_2_ gradients were analysed using real-time PCR assays targeting *Bacteria* using the primers 341F (5’-CCTACGGGAGGCAGCAG-3’) and 534R (5’-ATTACCGCGGCTGCTGGCA-3’) (see Supporting information for details).

### Bioinformatic processing of 16S rRNA gene amplicon data and statistics

The bioinformatic processing of 16S rRNA gene amplicon data was performed as previously described^53^. Briefly, read pairs were first assembled as contigs using QIIME’s join_paired_ends.py script^55^. Sequences were then screened for chimeras and clustered into species-level OTUs at 99% sequence similarity using the UPARSE pipeline (version 8.0.1623)^56^. This involved dereplication of sequences, removal of singletons, and determination of OTU centroids using a 1% radius. OTU abundances were determined by mapping filtered sequences to OTU centroids using 99% identity. OTU sequences were subsequently classified using the Naïve Bayesian classifier^57^, as implemented in mothur (version 1.33.0)^58^. A confidence cutoff of 50% was used for the classification with the SILVA 119 SSU NR99 database as taxonomic reference^59^.

The IMNGS webserver^60^ was used to identify the presence and relative abundances of sequences related to OTUs representative of the ^13^C-labelled taxa identified by SIP, in different publically available Short Read Archive (SRA) datasets. Sequence identity cut-offs of ≥95% for *Shewanella, Ca*. Izemoplasma and *Fusibacteraceae* OTUs and ≥97% identity to *Lutibacter* and *Nitrincolaceae* OTUs were used when querying the SRAs via IMNGS (Supp. Fig. 7). A minimum overlap of 100 bp was required between query and database sequences.

### Statistical analyses of 16S rRNA gene amplicon sequencing data

All statistical analyses of 16S rRNA gene amplicon data were performed using the Rhea package (version 1.0.1-5)^61^ as implemented in the R software environment (version 1.1.383). To detect OTUs enriched in relative abundances in the dense fractions of CsCl_2_ density gradients from ^13^C versus ^12^C treatments, the ‘Serial Group Comparisons’ pipeline of Rhea was applied. Default parameters were used except for the following: abundance_cutoff <- 0.001, prevalence_cutoff <- 0.1; max_median_cutoff <-0.001; ReplaceZero = “NO”. This pipeline used the Wilcoxon Signed Rank Sum Test for matched OTUs to test for significant differences in OTU relative abundances between two treatments, at each time point. Relative abundance data of OTUs from at least three selected heavy fractions (ranging from 1.726-1.742 g ml^-1^) from ^13^C-amended versus corresponding unamended (^12^C) control treatments, for day 4, 10 and 13, were used as input (Supp. Table 5). Data values obtained from fractions of >1.742 g ml^-1^ were excluded, since preliminary examination of plots of OTU relative abundances across gradients revealed very few taxa were labelled to such an extent that they increased in relative abundances in those fractions. For day 4, a ‘pseudo-replicate’ consisting of an average value from the available two heavy fractions was produced to result in triplicates and thus facilitate the statistical analysis (Supp. Table 5). To identify OTUs that were enriched in relative abundances in the different microcosm treatments versus the no-substrate controls, for each time point, the ‘Serial Group Comparisons’ pipeline of Rhea was applied as described above, using data from triplicate microcosms. Only OTUs that were significantly enriched in relative abundances at two or more time points, in each treatment series, were reported.

For beta diversity analysis of microbial communities in microcosms, principal coordinate analysis (PCoA) was performed using a Bray-Curtis dissimilarity matrix (including relative abundances). The plots were constructed with the ‘vegan’ package (version 2.5-3) in R. To avoid biases related to differences among library depths, sequencing libraries were all subsampled to a number of reads smaller than the smallest library (1620 reads).

### Metagenomic library preparation, sequencing and analyses

Metagenomic libraries were prepared for sequencing and indexed using the Nextera XT DNA Library Preparation Kit (Illumina, USA), following the manufacturer’s instructions. Five samples were selected to facilitate differential coverage binning, i.e., i) the initial sediment slurry, ii) day 4, ^12^C-DNA treatment, iii) day 13, ^13^C-DNA treatment, iv) day 24, ^12^C-DNA treatment, and v) day 31, ^13^C-DNA treatment. DNA libraries were purified using magnetic beads via the Agencourt AMPure XP kit (Beckman-Coulter), and sequenced by the Vienna BioCenter Core Facilities GmbH (VBCF) on one lane of an Illumina HiSeq 2500 instrument using HiSeq V4 chemistry with 125 bp paired-end mode.

Raw sequences were trimmed to 100 bp, and each sample dataset was assembled separately using IDBA-UD (version, 1.1.1) using default parameters^62^. Assembled contigs >1000 bp were automatically binned into genome bins based on a combination of nucleotide coding frequencies and sequence assembly coverage using MetaBat2 using each present binning strategy (version 2.12.1)^63^, MaxBin2 (version 2.2.4)^64^ and CONCOCT (version 0.4.1)^65^. Coverage profiles for binning were acquired by mapping trimmed reads to assemblies using BWA^66^ and SAMtools^67^. Additionally, the sample from the day 31 ^13^C-DNA treatment was also assembled using MetaSpades (version 3.11.1)^68^ and additionally binned with MetaBat, MaxBin and CONCOCT. Genome bins from each binning tool and from each respective assembly were aggregated using DasTool (version 1.1.0)^69^. All genome bins were finally dereplicated using dRep (version 1.4.3)^70^, with the following options: all genomes were dereplicated using an average nucleotide identity of ≥98% was used in the secondary ANI comparison, and all genomes that were >50% complete and <10% contamination were kept. The completeness and degree of potential contamination of the genome bins were evaluated by CheckM (version 1.0.7)^71^. Gene calling and initial genome annotations were performed via RAST^72^. Putative functions of predicted proteins were also checked via BLASTP^73^ against the NCBI-nr database and evaluated in relation to relevant literature. Predictions of subcellular locations of predicted proteins were performed using PSORTb (version 3.0)^74^, with settings appropriate for the predicted cell wall types of the respective organism. Biochemical inferences were primarily based on the MetaCyc database^75^. Gene synteny depictions were produced using EasyFig^76^.

### Phylogenetic and average nucleotide identity analyses

All phylogenetic analyses of 16S rRNA gene sequences were performed in ARB (version 6.0.6) using maximum likelihood algorithms^77^ and the SILVA 132 SSU NR99 database^59^. Query sequences were first aligned to the SILVA alignment with the SINA aligner^78^, then sequences were inserted into the global reference tree using the parsimony option. A selection of near-full length sequences of close relatives and cultivated relatives of the query sequences were then selected and used to construct trees *de novo* using RaxML^79^, fastDNAmL^80^ and PhyML^81^ algorithms as implemented in ARB with default settings and the ARB bacterial ‘Positional variability by parsimony’ filter. A consensus tree was then created with those three trees and short amplicon sequences were inserted into the consensus trees via the parsimony option of ARB.

Phylogenomic analyses were conducted by using an alignment of concatenated protein sequences derived from 43 single copy marker genes retrieved from the CheckM analyses. A tree was first constructed using these protein alignments derived from the query genomes and 8203 representative genomes downloaded from the NCBI database (as of July 2018). The tree was constructed using default parameters using FastTree (version 2.1.10)^82^. Manually selected reference sequences that were taxonomically informative for query sequences were obtained, and a second smaller tree was reconstructed using FastTree using only the selected reference and query sequences.

Average nucleotide identity (ANI) for comparison of genome relatedness were determined using JSpeciesWS server^83^, based on BLAST (‘ANIb’).

### Taxonomic names

In addition to 16S rRNA-based SILVA taxonomies, we used the newly proposed names of the Genome Taxonomy Database (GTDB) (version 0.1.3) that is based on genome phylogeny^46^.

### Chemical measurements

Carbon isotope compositions of CO_2_ (δ^13^C values in permille relative to VPD) were analysed by a headspace gas sampler (Gas-Bench II, Thermo Fisher, Bremen, Germany) coupled to an isotope ratio mass spectrometer (Delta V Advantage, Thermo Fisher, Bremen, Germany). CO_2_ reference gas was calibrated using ISO-TOP gas standards (Air Liquide) with a certified ^13^C concentrations.

For total sediment iron and manganese, inductively coupled plasma optical emission spectrometry (ICP-OES) measurement was applied on a Perkin Elmer 5300 DV (Pekin Elmer Inc., Waltham, MA, USA) after total fusion of 0.1 gram oven dried sediment samples (1 g original sample dried 3 h at 105°C, loss of ignition 6 h at 600°C, followed by 6 h at 1,000°C, LOI approx.. 80%) with 0.9 g di-lithium tetraboarate (Spectromelt A10, Merck, Darmstadt, Germany) using a Linn Lifumat, 9 min. at 1,050 °C (Linn High Therm Gmbh, Eschenfelden, Germany). The fusion product was dissolved in 50 ml of 5 molar HNO_3_ (Normapur, VWR, Germany, double sub-boiled, Berghof BSB-939-IR) with 150 ml deionized water (MilliQ, Merck-Millipore, Darmstadt, Germany), and further diluted to a total volume of 250 ml. External linear matrix matched calibration was applied using single element manganese and iron standards (Merck-Millipore, Darmstadt, Germany), including quality control with LKSD1 and LKSD 4 certified lake sediment standards (Natural Resources Canada, Ottawa, Canada).

## Supporting information

Supp. Table 1 - Treatment summary

Supp. Table 2 - Genome statistics

Supp. Table 3 - List of key annotations for DNA degradation in MAG

Supp. Table 4 - De novo syntheses of nucleotides

Supp. Table 5 - Gradient fractions used for sequencing and statistics

Supp. Fig. 1 - Headspace CO2 isotopic compositions

Supp. Fig. 2 - Halobacterium relative abundances in microcosms

Supp. Fig. 3 - Microcosm microbial community bar graphs

Supp. Fig. 4 - PCoA plot of microbial communities in microcosms

Supp. Fig. 5 - qPCR of SIP fractions

Supp. Fig. 6 - Relative abundances of taxa in SIP gradients

Supp. Fig. 7 - Global distributions of DNA-degrader OTUs

Supp. Fig. 8 - Microcosms summary - OTUs enriched at 2 time points

Supp. Fig. 9 - Xanthine dehydrogenase operon synteny

Supp. Fig. 10 - Halobacterium in SIP gradients

Supp. Fig. 11 - BLASTP domain result

## Data availability

All sequence data was deposited under Genbank Bioproject PRJNA510104. PCR-derived 16S rRNA gene amplicon sequence data is available under accessions SAMN10603326-SAMN10603488. Metagenomic sequence read data is available under accessions SAMN10594394-SAMN10594398. Metagenome-assembled genomes are under processing and will be available under accessions XXXX.

## Acknowledgements

This work was supported by the Austrian Science Fund (FWF) grants P25111-B22 and P29246-B29 to A.L. and K.W., respectively. We thank the Vienna BioCenter Core Facilities for metagenomic sequencing. We thank Prof. Bernhard Schink and Prof. Aharon Oren for their recommendations for nomenclature proposed in this study. We thank Dr. Angela Witte for kindly providing us with the *Halobacterium salinarum* strain and Fátima C. Pereira for performing the PCoA analysis. For sampling, funding from ArcticNet and assistance from the Canadian Coast Guard, captain and crew of the CCGS *Amundsen*, and chief scientist are gratefully acknowledged.

## Author contributions

K.W. and A.L. conceived and designed the study. A.N. and C.R.J.H were responsible for sample collection and processing. K.W. and C.P. performed the incubation experiments and molecular analyses. M.W. and A.R. performed gas chromatography analyses. T.H. performed sediment iron and manganese analyses. K.W., C.P. and C.W.H performed bioinformatic processing and analyses. K.W. analysed and interpreted the data. K.W. and A.L. wrote the manuscript with contributions from all authors.

## Materials & Correspondence

Requests for materials and/or correspondence can be made to Kenneth Wasmund and/or Alexander Loy

