## Supporting Information - TEXT for "DNA-foraging bacteria in the seafloor"

### Supporting Materials and Methods

#### Artificial seawater

Artificial seawater was prepared with 20 g $NaCl_2$, 3 g $MgCl_2 \cdot 6H_2O$, 0.15 g $CaCl_2 \cdot 2H_2O$, 0.3 g KCl, 0.25 g $NH_4Cl$, 0.2 g $KH_4PO_4$ and 3.9 g $Na_2SO_4$ per liter of $ddH_20$. The pH was adjusted 7.6. The solution was autoclaved and then transferred into an anoxic glove-box while warm. Resazurin was added to a final concentration of 0.5 mg $l^{-1}$. 1,4-Dithiothreitol (DTT) was also added as reducing agent to 250 µM final. The solutions was further deoxygenated within the anoxic tent for 3 days with the lid of the Schott bottles slightly opened. Solutions were then sealed with rubber stoppers and cooled to 4°C before the experiment was set-up.

#### Growth of *Halobacterium salinarum*

*Halobacterium salinarum* was grown in a medium with 1.25 g NaCl, 0.1 g $MgSO_4$, 0.04 g KCl, 0.006 g sodium citrate, and 'carbon mix' 0.05 g, per 5 ml $ddH_2O$. 'Carbon mixes' were either i) ISOGRO®-$^{13}C$ Powder-Growth Medium (99 atom % $^{13}C$) (Sigma-Aldrich), or ii) $^{12}C$ yeast extract (Oxoid). Cultures were grown at 37°C for up to 3 days.

#### Preparation of $^{13}C$- and $^{12}C$-labelled DNA for use as substrate

*Halobacterium salinarum* that was grown on either i) ISOGRO®-$^{13}C$ Powder-Growth Medium (99 atom % $^{13}C$) (Sigma-Aldrich) for three generations, or ii) $^{12}C$ yeast extract, was extracted using a CTAB-phenol method as described for sediment DNA (see below), except cells were pelleted and supernatant removed before DNA extraction. DNA extracts from the *Halobacterium* were washed with a Microcon YM-10 (Millipore) device to reduce any possible residual low molecular solutes less than approximately 10 kDa. The extracts had 260/280 ratios averaging around 2.0 as determined by spectrophotometry using a Nanodrop-1000 device (Nanodrop). The highly gelatinous consistency of the purified extract further indicated that DNA was the major organic compound in the extract. The $^{13}C$ enriched nature of the *Halobacterium* DNA extract was demonstrated by examination of *Halobacterium* 16S rRNA gene abundances in SIP density gradients, which showed these sequences dominating in the most dense ('heaviest') fractions retrieved (Supp. Fig. 10).

#### Extraction of genomic DNA

DNA was extracted from sediments and *Halobacterium salinarum* using a combination of bead beating, cetyltrimethylammonium bromide-containing buffer and phenol-chloroform extraction. Briefly, sediment samples or pelleted cells were suspended in 400 µl of extraction buffer (100 mM Tris-HCl [pH 8.0], 100 mM sodium EDTA [pH 8.0], 100 mM sodium phosphate [pH 8.0], 1.5 M NaCl, 1% [wt/vol] cetyltrimethylammonium bromide). An additional 55 µl of sodium dodecyl sulfate (20%, wt/vol) was also added. This was transferred to a Lysing Matrix E 2 ml tube (MP Biomedicals) and the samples were subject to bead beating at 'speed 6' for 2 rounds of 30 s using a FastPrep®-24 bead beating instrument (MP Biomedicals), with cooling on ice between. The supernatant was decanted into a clean tube after centrifugation for 10 min at 6,000 g. An equal volume of phenol:chloroform (containing 4% [vol/vol] isoamyl alcohol) was then added and the samples were mixed by inversion, and then centrifuged at 16,000 g for 10 min at 25°C. The aqueous phase was collected to a new tube and precipitated with 0.6 volume of isopropanol at 4°C for 30 mins. The precipitate was pelleted by centrifugation at 16,000 g for 30 min at 4°C, the supernatant removed carefully, and then washed with 70% (vol/vol) ethanol and air dried for 5 mins. The DNA was finally resuspended in 100 µl $ddH_20$.

#### PCR amplification of 16S rRNA genes for amplicon sequencing

PCR reactions for the first-step PCR (total volume of 12.5 µl) contained 7.45 µl ddH20, 12.5 µl of 10× Taq Buffer (Thermo Scientific), 250 µM dNTP mix, 1.5 mM MgCl2, 0.2 µM of each primer, 0.05 µl Thermo Scientific Taq DNA Polymerase (Thermo Scientific) and 0.5 µl of DNA template (1:100 dilutions of crude extracts). PCR conditions for the first-step PCR were: 95°C for 2 min, followed by 30 cycles of 95°C for 30 s, 52°C for 30 s and 72°C for 30s, followed by a final extention of 72°C for 2min. For the second-step barcoding PCRs, 1 µl of the first-step PCR was added to 25 µl PCRs containing the same concentrations of reagents detailed as for the first-step PCR, except the barcoded primer was 0.4 µM in each reaction. PCR conditions for the second-step PCR were the same as for the first-step PCR, except a total of 15 cycles were performed.

#### Quantitative real-time PCR assays

Real-time PCR reactions (10 µl total) contained 5 µl iQ™ SYBR® Green Supermix (BioRad, Austria), 0.5 µM of each primer 341F (CCTACGGGAGGCAGCAG) and 534R (ATTACCGCGGCTGCTGGCA), 3 µl ddH20 and 1 µl of DNA template (1:100 dilutions of crude extracts). PCR conditions were: 95°C for 3 min, followed by 40 cycles of 95°C for 30 s, 60°C for 30 s and 72°C for 30s, followed by a final extention of 72°C for 2 min. Standards included a 16S rRNA gene derived from an in-house culture of *Desulfosporosinus acidiphilus*, which was cloned into a pPCR4-TOPO vector (Invitrogen, USA) and reamplified by PCR using M13 primers, and then PCR purified and diluted to a range from $10^9$ to $10^3$ copies per microliter. All real-time PCR reactions were performed in triplicate. Real-time PCR was performed with a CFX96 Touch™ Real-Time PCR Detection System (version 3.1) (Bio-Rad). A melting curve analysis was also performed after the final extension using default instrument settings, whereby the temperature was increased from 65°C to 95°C.

### Supporting Results and Discussion

#### Microbial community structures and biogeochemical processes in microcosms

The overall bacterial community structures among microcosms shifted over the time course of the experiment, yet were similar between no-substrate control microcosms and microcosms supplemented with $^{12}$C- or $^{13}$C-DNA over time (Supp. Fig. 3 and 4). This indicated that the amounts of added DNA did not significantly alter the overall community dynamics. The communities were generally dominated by the deltaproteobacterial families *Geobacteraceae* and *Desulfuromonadaceae*, which averaged 15% and 11% over all time points, respectively. Other predominant taxa were from the gammaproteobacterial families *Shewanellaceae*, *Colwelliaceae*, *Nitrincolaceae* (GTDB, formerly *Oceanospirillaceae*), and the epsilonprotoebacterial family *Campylobacteraceae*, which averaged 3.5%, 4.7%, 14.5% and 7.5% over all time points, respectively. Members of the families *Flavobacteriaceae*, *Ca*. Izemoplasmataceae (formerly *Tenericutes* family 'NB1-n'), and *Fusibacteraceae* (GTDB, formerly 'Clostridial Family-XII') increased in relative abundances towards the end of the incubations and made up to 11%, 5% and 14.5% of the sequences at day 31, respectively. The high relative abundances of *Desulfuromonadaceae*, *Geobacteraceae*, *Shewanellaceae*, *Colwelliaceae*, *Campylobacteraceae* and *Oceanospirillaceae* (GTDB name *Nitrincolaceae*) is conspicuously similar to the composition of taxa previously found to dominate in manganese-rich marine sediments from various locations[1].

Chemical analyses indicated that sulfate was not depleted over the time course of the incubations (results not shown) and no indications for sulfide production could be detected by smell in any of the microcosms at the end of the experiment. Measurements of total manganese and iron demonstrated high concentrations of these metals in the sediments, i.e., 6.25 and 40.4 g $kg^{-1}$ (dry weight), respectively. This indicated that metals were abundant potential electron acceptors for microorganisms during the anoxic incubations. Together with the high prevalence of typical metal-reducing taxa (see above), this indicated that the reduction of metals such as manganese and/or iron was likely the predominant terminal respiratory process during this experiment.

#### DNA-degrading taxa are globally distributed in marine sediments

To examine the global distributions of the five DNA-degrading bacterial genera/species identified in the SIP analyses, we examined the presence of 16S rRNA gene sequences closely related to OTUs that were determined to be labelled in the SIP analyses (Fig. 1) in publically available Short Read Archive (SRA) datasets. We identified 247 SRA samples from marine sediments that simultaneously contained sequences from all of the five genera/species. The relative abundances of these taxa in representative samples of each of the sites (n=18) showed that these taxa are globally distributed (Supp. Fig. 7). The samples included diverse marine ecosystems, from both coastal to deep-sea environments, as well as polar and tropical marine environments. They also spanned distant locations, from both Arctic and Antarctic regions, as well as from European, Asian and North and South American continental margins. At the most, these five taxa collectively represented 10% of some microbial communities in methane seep associated sediments off the coast of Oregon, USA. Sequences related to *Ca*. Izemoplasma OTU 23 were found up to 2% in river delta sediments of the Adriatic Sea, Italy. Sequences related to *Nitrincolaceae* OTU 4994 were only found at a maximum of 0.3% relative abundance.

#### Description of relatedness of MAGs to known organisms and genomes

Here we provide a description of the phylogenetic affiliations and novelty of MAGs recovered in this study revealed by phylogenomic analysis (Fig. 2), genome based average nucleotide identity (ANI) analyses (Supp. Table 2), as well as in relation to 16S rRNA genes recovered from the same samples. Taxonomic cut-offs based on genome ANI[2] and 16S rRNA sequence identities [3] were considered. The *Lutibacter* MAG BB-2 was phylogenetically affiliated with the genus *Lutibacter*, yet could be considered a novel species based on an ANI of 78% to *Lutibacter flavis*, and the fact that no recovered 16S rRNA genes had >98% identity to any *Lutibacter* spp.. The *Nitrincolaceae* MAG BB-4 was phylogenetically related to, yet rather distinct from *Motiliproteus* sp. MSK22_1. Further, based on low ANI (69%) from very low amounts of aligned regions (31%), and the fact that all *Nitrincolaceae* 16S rRNA gene OTUs were <94% related to any described species, the *Nitrincolaceae* MAG BB-4 may thus represent a novel genus within the family *Nitrincolaceae*. The *Shewanella* MAG BB-1 was phylogenetically affiliated within the genus *Shewanella* and most related to *Shewanella sediminis.* Nevertheless, it probably constituted a novel species based on 86.5% ANI from 62% of aligned genome to *Shewanella sediminis*. The *Fusibacteraceae* MAG BB-3 was broadly related to *Fusibacter* sp. 3D3 based of phylogenomic analysis, had only 65.2% ANI from only 16% aligned, and together with the fact that 16S rRNA gene OTUs from this family had <94% identity to any cultured species, it may thus represent a novel genus. The *Ca.* Izemoplasma acidinucleici MAG was most related to yet clearly distinct from *Ca.* Izemoplasma sp. HR1, with 70.5% genome ANI from only 42.9% aligned. The assembled 16S rRNA gene of *Ca.* Izemoplasma MAG had only 94% sequence identity to *Ca.* Izemoplasma sp. HR1. This MAG therefore represents a novel species for which we propose the name *Ca.* Izemoplasma acidinucleici.

#### Genes for *de novo* biosynthesis of nucleotides in *Ca*. Izemoplasmataceae genomes

Among *Ca*. Izemoplasma, all genes except one required for *de novo* syntheses of purines could be identified (Supp. Table 4). Only a nucleoside diphosphate kinase required for phosphorylation of GDP to GTP was not encoded. Nevertheless, this pathway must be performed by an alternative means otherwise the organisms would not be viable. Instead, it may be performed by a predicted multi-functional adenylate kinase, as was previously predicted to perform this function in related phytoplasmas[4]. Similarly, all genes required for complete *de novo* pyrimidine biosyntheses from glutamine and bicarbonate could be identified among *Ca*. Izemoplasma except for a canonical carbamoyl phosphate synthetase, which was only present in *Ca*. Izemoplasma acidinucleici (Supp. Table 4). Instead, we identified genes for carbamate kinases in most *Ca*. Izemoplasma that may instead catalyze this step, as was shown previously for various archaea[5].

#### Large protein with DNA-binding and nuclease domains

Encoded within the ‘DNA-degradation loci’ of various *Ca*. Izemoplasmatales (Fig. 5) was a ‘Large protein with nucleic acid-binding residues, C-terminal endonuclease I domain and ‘Lamin Tail Domain’. These domains were detected by BLASTP against the NCBI's Conserved Domain Database (CDD)[6] (Supp. Fig. 11). Details regarding these domains include:

- ‘Generic binding surface’ domain:
    - Mostly consists of nucleic acid substrates in most OB-fold complexes[6].
- YhcR_OBF_like domainv (RPA_2b):
    - Subfamily of OB-fold domains *Bacillus subtilis* YhcR. The YhcR consists of a sugar-nonspecific nuclease[6].
- ‘Lamin Tail Domain’ (LTD):
    - In some secreted or periplasmic proteins, LTDs are associated with other substrate-binding domains such as those for oligosaccharides. These associations indicate possible roles for LTDs to act as tethering proteins to membranes or associated protein[7].

We hypothesize that this large protein could function to hold polymeric DNA in close proximity to the cell during digestion, since it contains the domains necessary to do so. In that way, DNA could be degraded in concert with the other secreted nucleases close to the cell, thereby minimizing diffusion of liberated sub-components and enable efficient uptake by the associated transporters.
