## Supplementary figures and images for "DNA-foraging bacteria in the seafloor"

### Supp. Fig. 1 - Headspace CO2 isotopic compositions

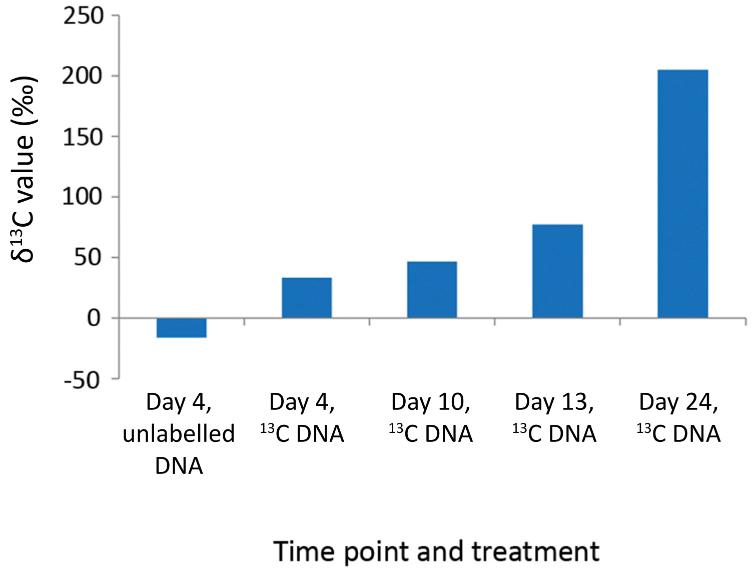


**Supp. Fig. 1.** Isotopic composition of $CO_2$ from the headspace of representative microcosms.
