## Supplementary material for "DNA-foraging bacteria in the seafloor": Supp. Fig. 2 - Halobacterium relative abundances in microcosms

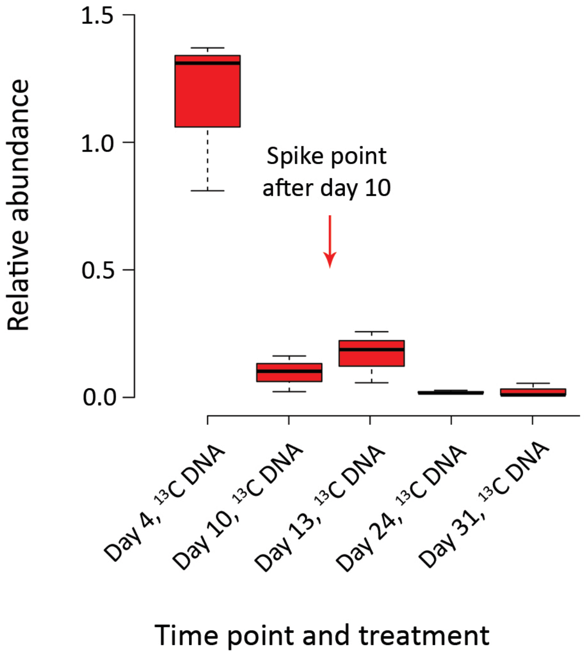


**Supp. Fig. 2.** Relative abundances of 16S rRNA genes of the archaeon *Halobacterium salinarum*, which was added to the microcosms as substrate. It should be noted that relative abundances were not determined at Day 0 because the *Halobacterium* DNA was supplemented after sub-sampling. 'Spike point' denotes the day at which additional *Halobacterium* DNA was added to the microcosms after sub-sampling (see Materials and Methods).
