## Supplementary material for "DNA-foraging bacteria in the seafloor": Supp. Fig. 3 - Microcosm microbial community bar graphs

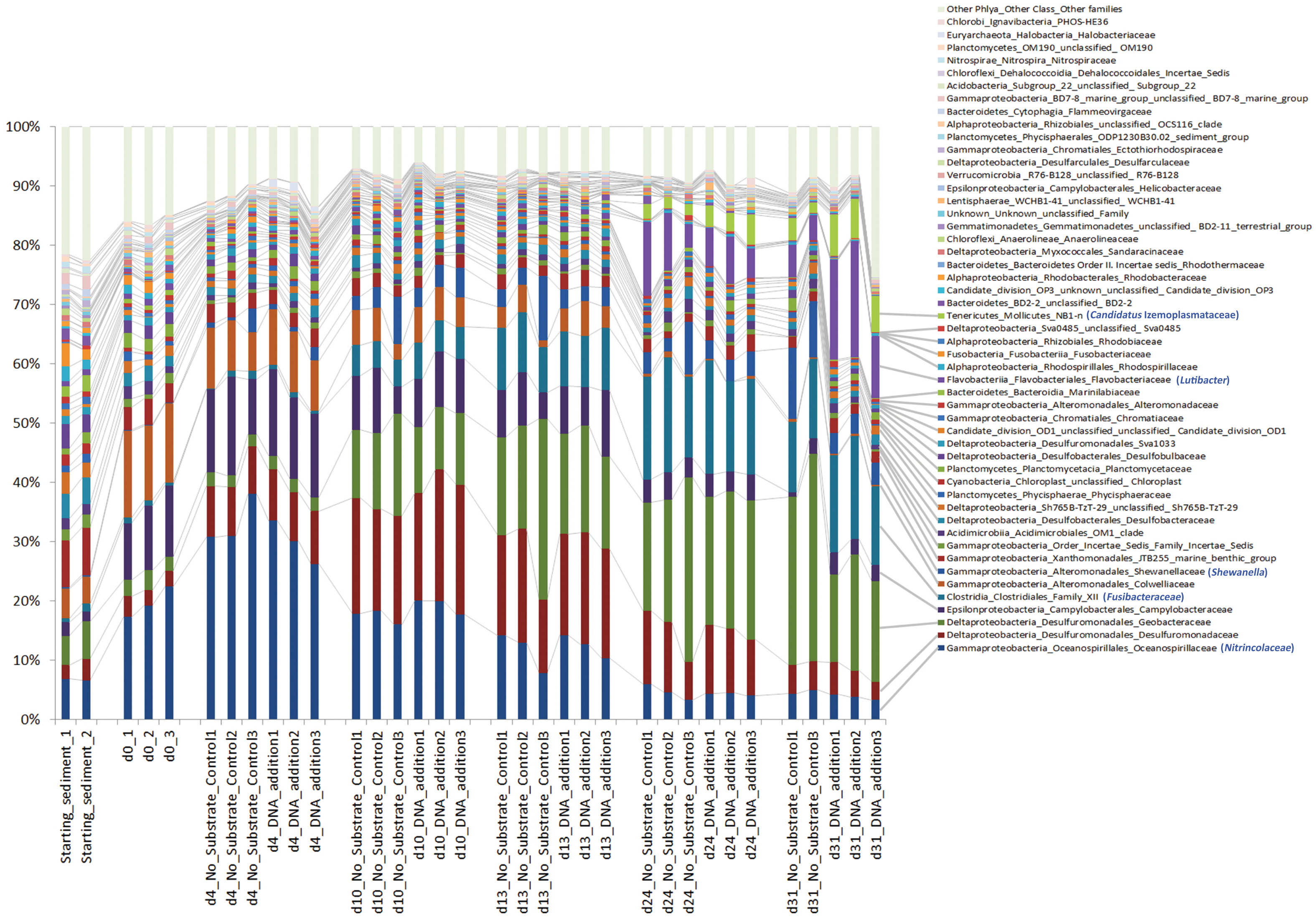


**Supp. Fig. 3.** Microbial community structures among starting sediments and microcosms throughout the incubation series. The graph is based on 16S rRNA gene amplicon sequencing data. Relative abundances of family-level classifications are presented, with 'Phyla_Order_Family' taxonomic strings presented for all non-proteobacterial lineages, and 'Class_Order_Family' taxonomic strings for all proteobacterial lineages. SILVA 16S rRNA taxonomic names are presented for all taxa, and Genome Taxonomy Database (GTDB ) taxonomic names for taxa determined to be $^{13}$C-labelled in SIP (Fig. 1) are given in blue parentheses. The top 50 most abundant families are presented, and all others are combined as 'Other Families'. Only $^{12}$C-DNA supplemented treatments and 'No Substrate Controls' are shown for clarity. NMDS plots comparing Beta-diversity among $^{12}$C-DNA and $^{13}$C supplemented treatments showed little differences and are summarized in Supp. Fig. 4.
