## Supplementary material for "DNA-foraging bacteria in the seafloor": Supp. Fig. 4 - PCoA plot of microbial communities in microcosms

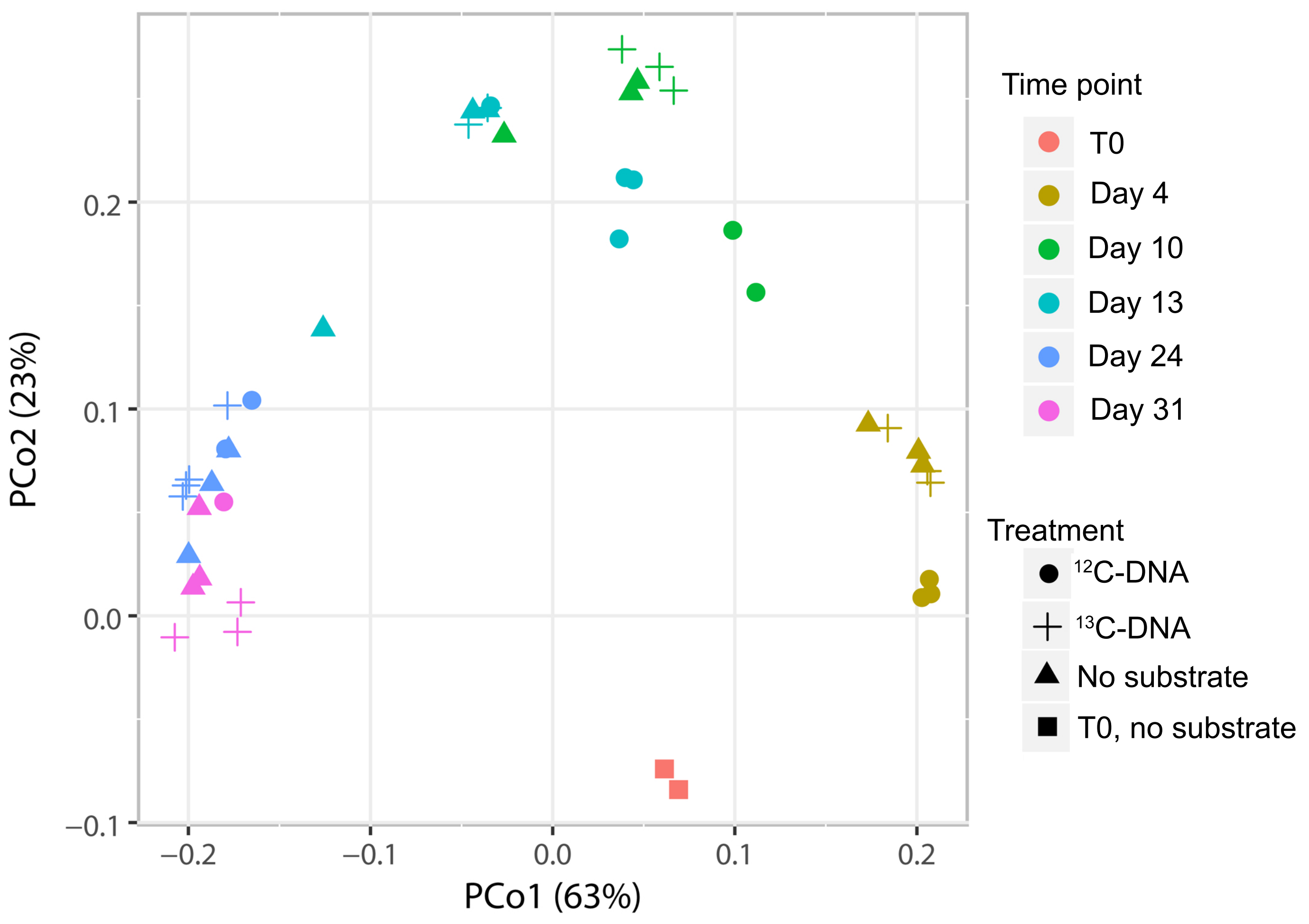


**Supp. Fig. 4.** PCoA plot of microbial community differences among microcosms supplemented with DNA versus controls, over time.
