## Supplementary material for "DNA-foraging bacteria in the seafloor": Supp. Fig. 5 - qPCR of SIP fractions

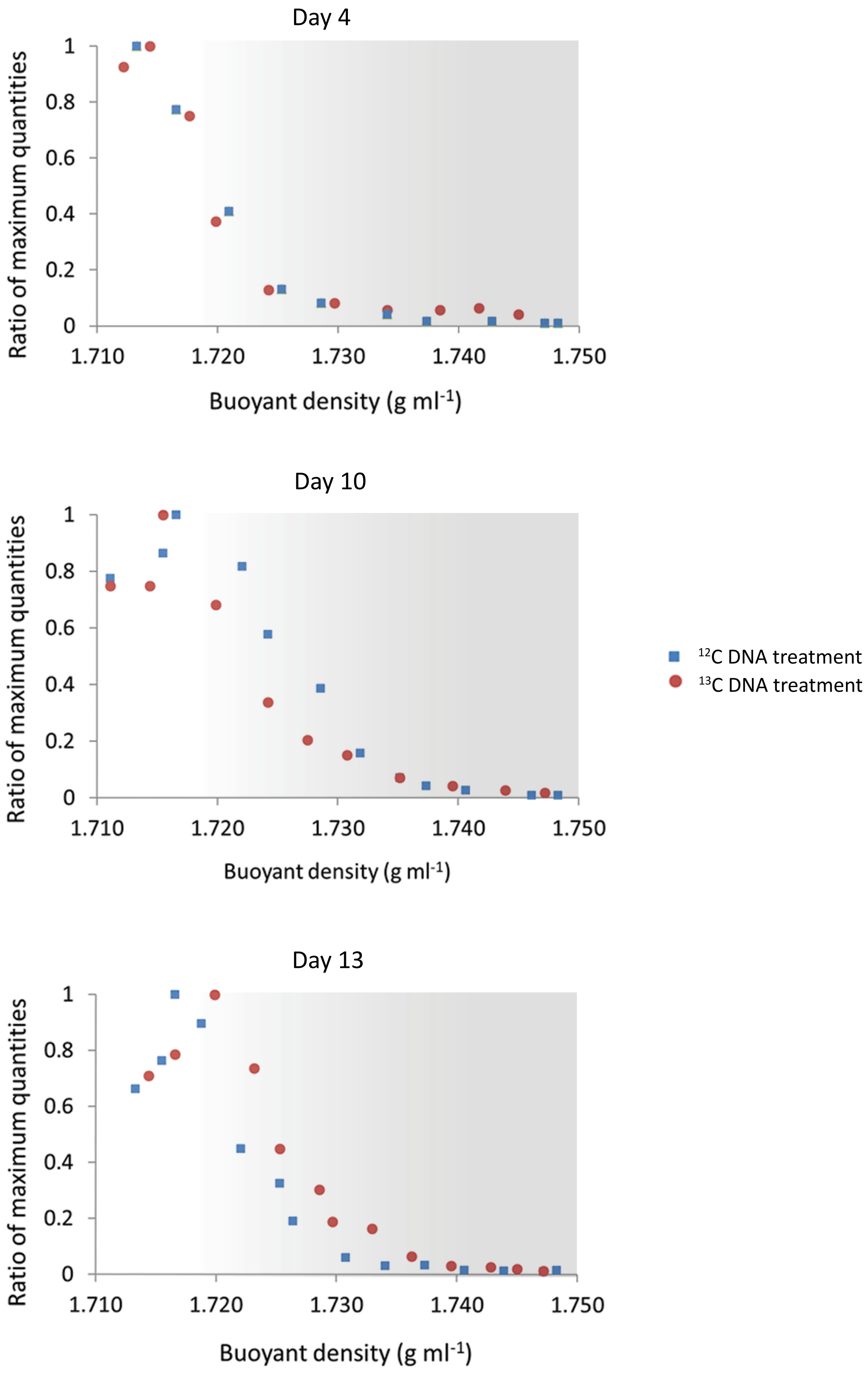


**Supp. Fig. 5.** Quantification of total bacterial 16S rRNA genes across buoyant density gradient fractions collected from treatments supplemented with $^{13}$C-DNA or $^{12}$C-DNA, as assessed by real-time PCR. Values presented are the ratio of maximum quantities detected in each series.
