## Supplementary material for "DNA-foraging bacteria in the seafloor": Supp. Fig. 6 - Relative abundances of taxa in SIP gradients

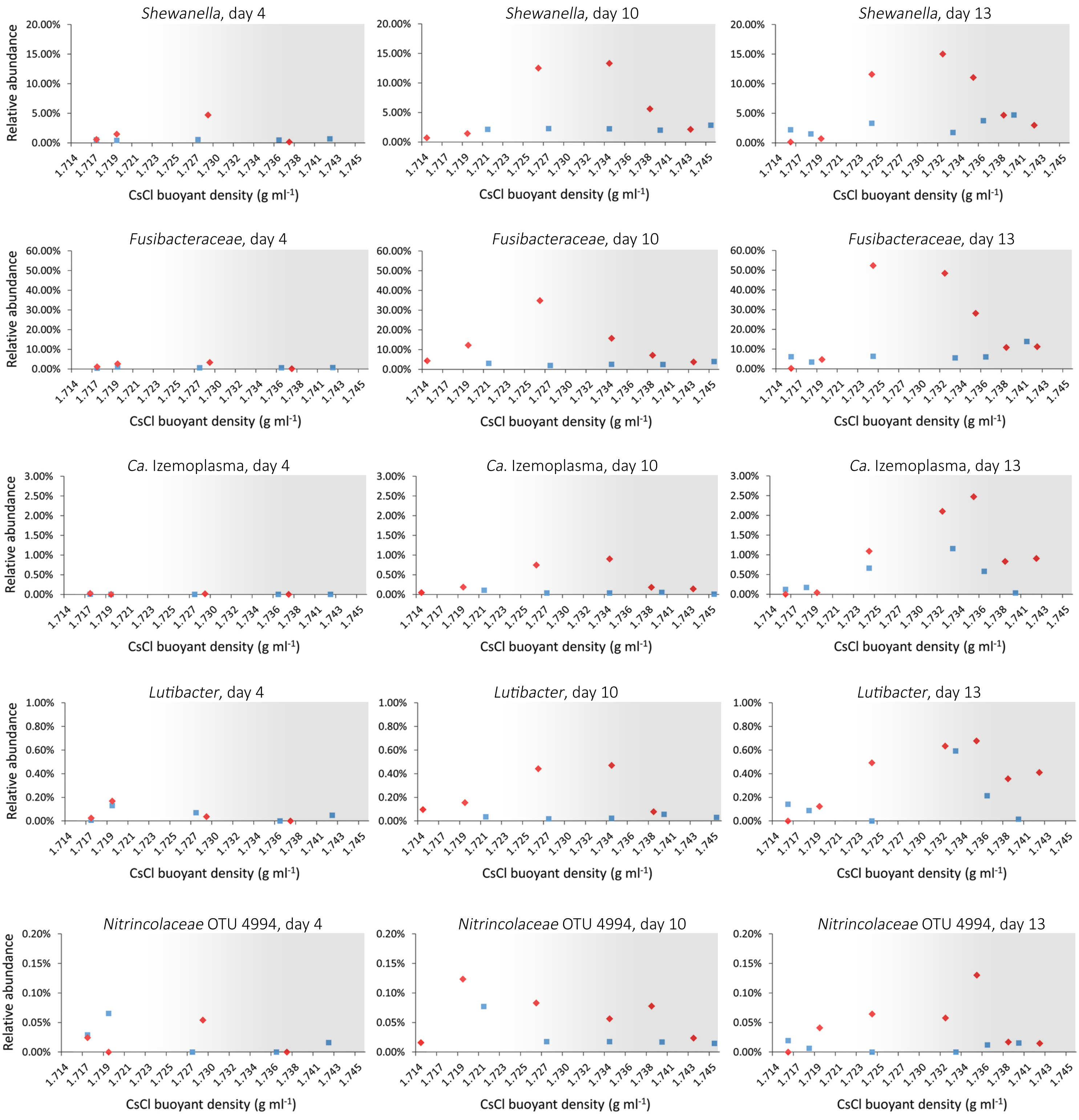


**Supp. Fig. 6.** Summed relative abundances of OTUs in taxonomic groups determined to be enriched in $^{13}C$ by amplicon sequencing of 16S rRNA genes across isopycicnic density gradient fractions. Relative abundances determined across $^{13}C$ and $^{12}C$ gradients are plotted in red diamonds and blue squares, respectively. Grey shaded areas indicate the 'heavy' ends of the gradients. For the family *Nitrincolaceae*, only the relative abundances of *Nitrincolaceae* OTU 4994 are presented, since this group did not show 'heavy' labelling patterns when OTUs from this group were summed together. The number of OTUs that had their relative abundances summed for the *Shewanella*, *Fusibacteraceae, Ca.* Izemoplasma *and Lutibacter* were 12, 39, 25 and 31, respectively.
