## Supplementary material for "DNA-foraging bacteria in the seafloor": Supp. Fig. 7 - Global distributions of DNA-degrader OTUs

**Supp. Fig. 7.** Global presence and relative abundances of 16S rRNA gene sequences that are related to the top OTUs identified as labelled in SIP analyses when $^{13}$C-DNA was added as substrate. 16S rRNA gene sequences were examined in publically available Short Read Archieve (SRA) datasets via the IMNGS webserver (see Materials and Methods). In total, 247 SRA samples from marine sediments that simultaneously contained sequences from all of the five taxa were identified, and the relative abundances of these taxa in representative samples of each of the sites (n=18) are presented. Five additional samples were identifed with high abundances of *Ca*. Izemoplasma OTU 23 related sequences and were also plotted. Sequence identity cut-offs of ≥95% for *Shewanella*, *Ca*. Izemoplasma and *Fusibacteraceae* OTUs and ≥97% identity to *Lutibacter* and *Nitrincolaceae* OTUs were used when querying the SRAs via IMNGS, in order to reflect the phylogenetic breadth of the labelling among related OTUs detected by SIP.

*Ca*. Izemoplasma OTU 23

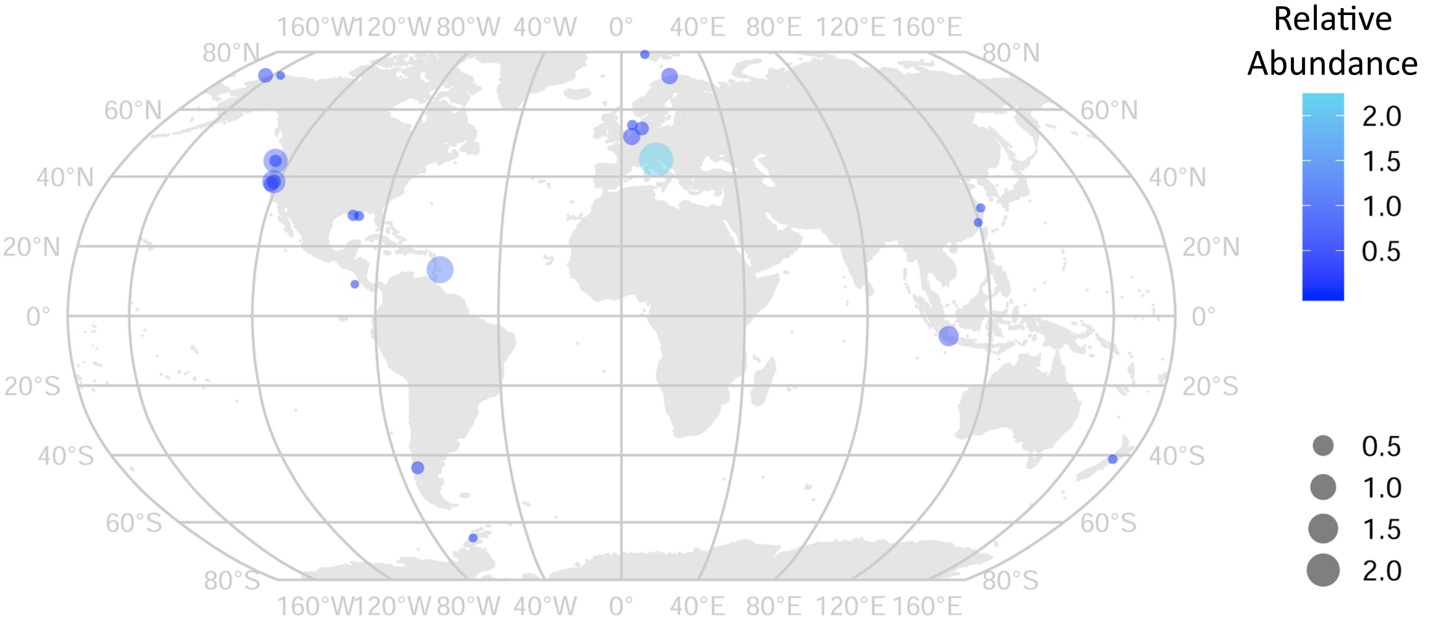


*Shewanella* OTU 12

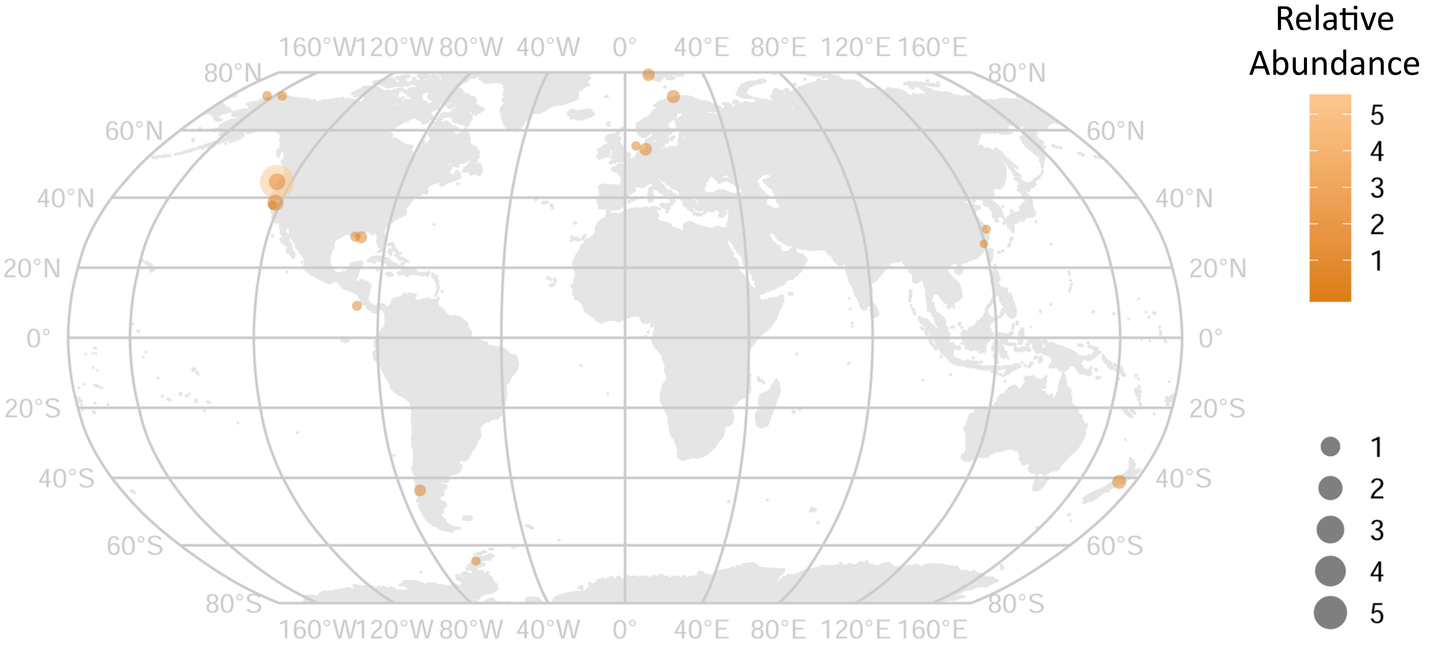

*Lutibacter* OTU 13

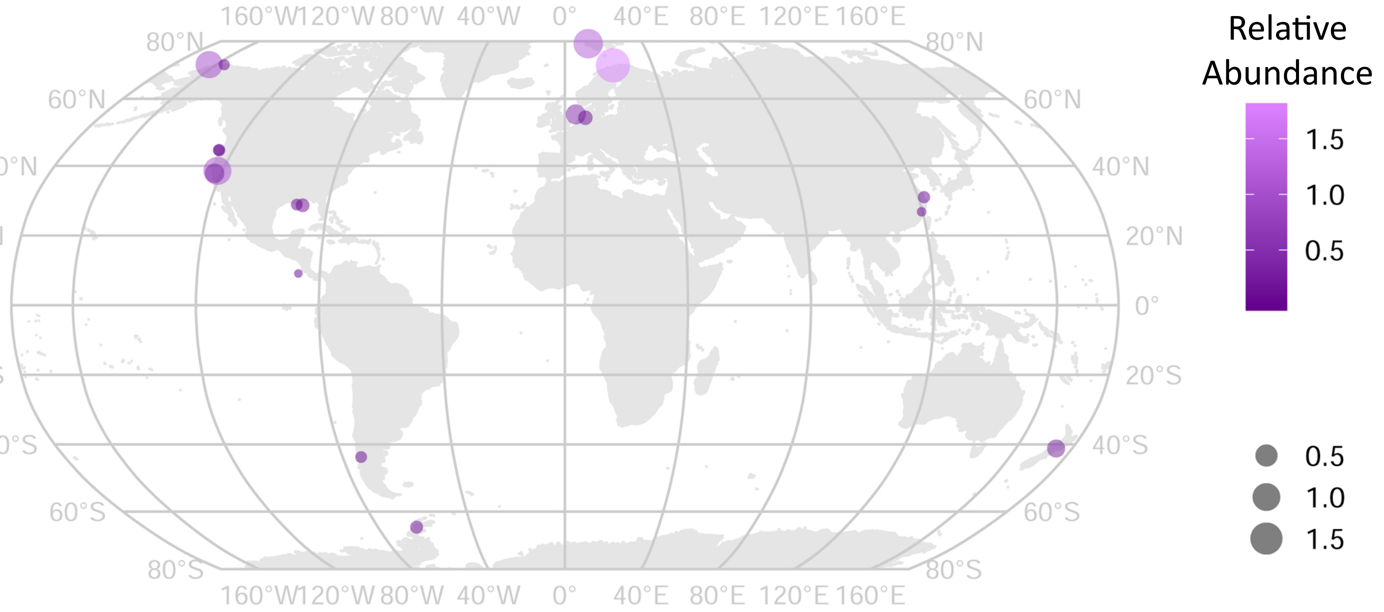


*Fusibacteraceae* OTU 6

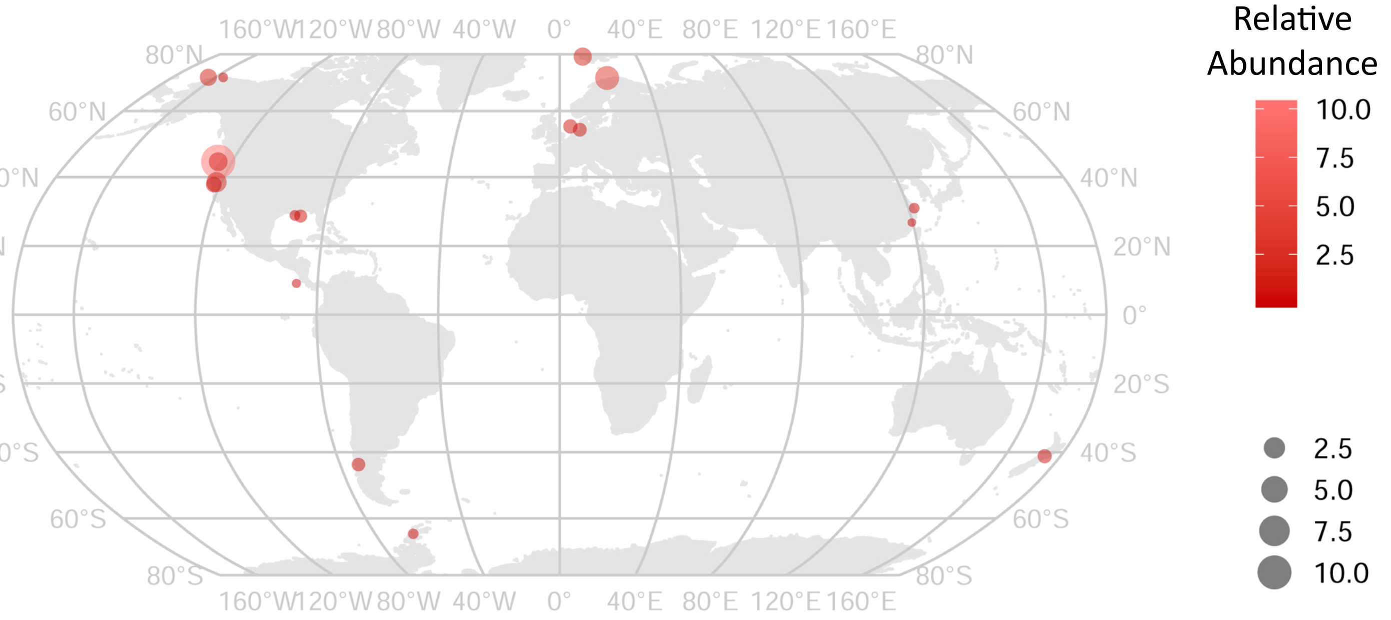


*Nitrincolaceae* OTU 4994

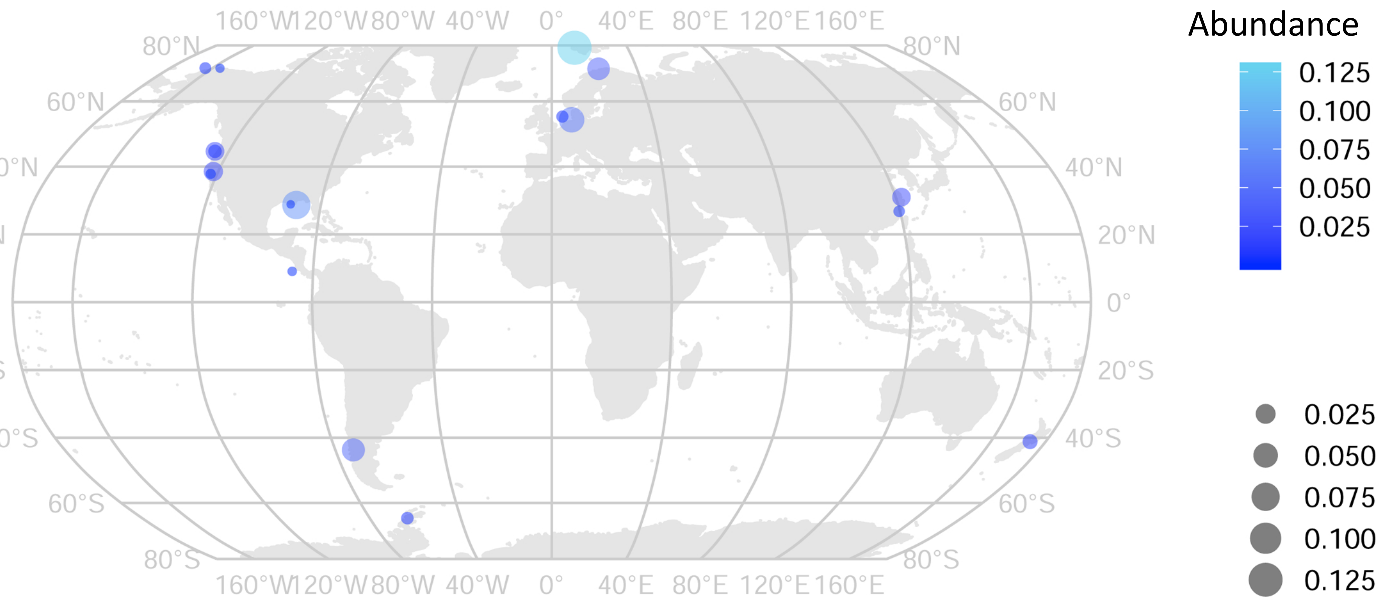
