## Supplementary material for "DNA-foraging bacteria in the seafloor": Supp. Fig. 8 - Microcosms summary - OTUs enriched at 2 time points

**Supp. Fig. 8**. Box plots of 16S rRNA gene relative abundances of all OTUs that were significantly enriched (p-value $<0.05$) in any treatment versus the same time-points corresponding no-substrate control, at $\geq 2$ time points. Red stars indicate which treatments and time points were significantly different from their corresponding no-substrate controls. The graphs are scaled to show significantly different treatments, i.e., high abundances may be cut-off. Treatments for nucleobases and nucleosides were only sequenced from the first three time points, i.e., days 4, 10 and 13. Taxonomic strings displayed = Domain;Phylum;Class;Order;Family;Genus; and are derived from the classifications against SILVA 16S rRNA gene database (see Materials and Methods). Further taxonomic information in relation to the Genome Taxonomic Database (GTDB) can be interpreted from Fig. 1.

**OTU 5:**

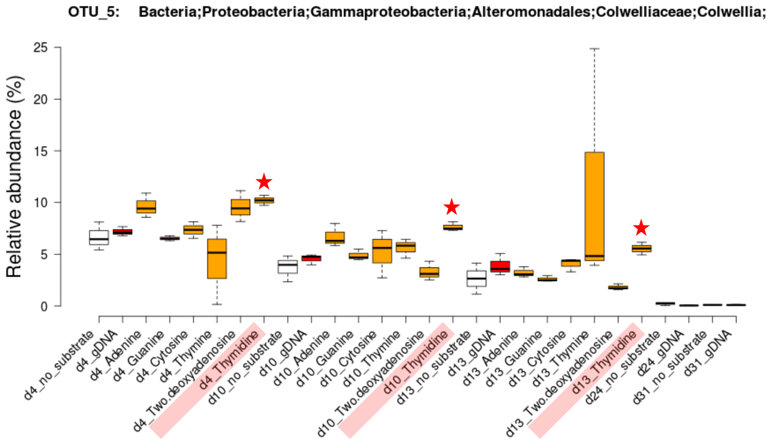


**OTU 6:**

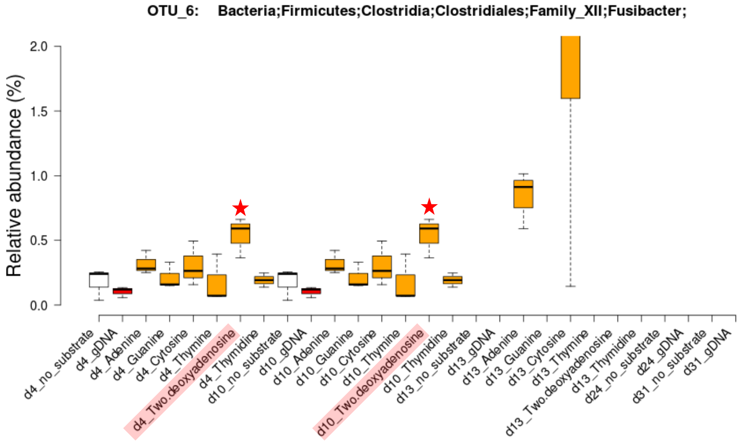


**OTU 7:**

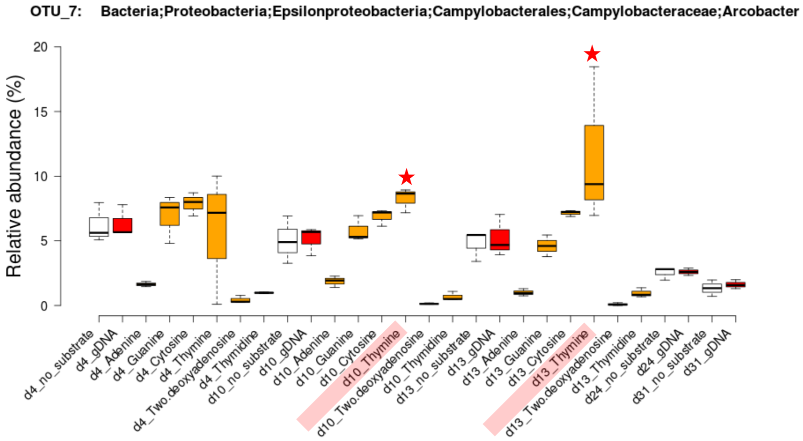


**OTU 8:**

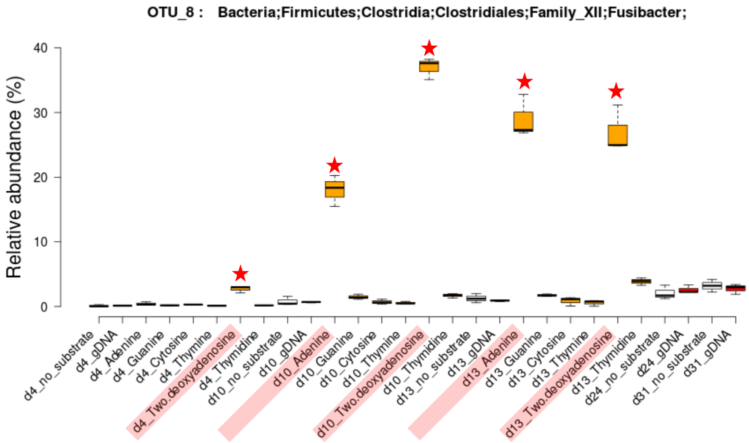

OTU 9:

OTU_9: Bacteria;Proteobacteria;Epsilonproteobacteria;Campylobacterales;Campylobacteraceae;Arcobacter

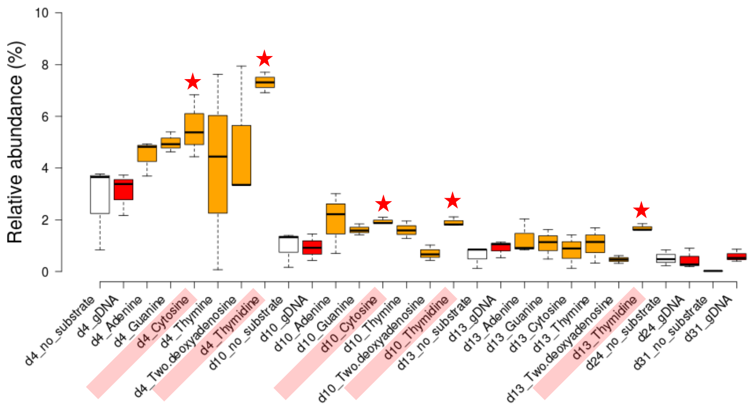


OTU 11:

OTU_11: Bacteria;Firmicutes;Clostridia;Clostridiales;Family_XII;Fusibacter;

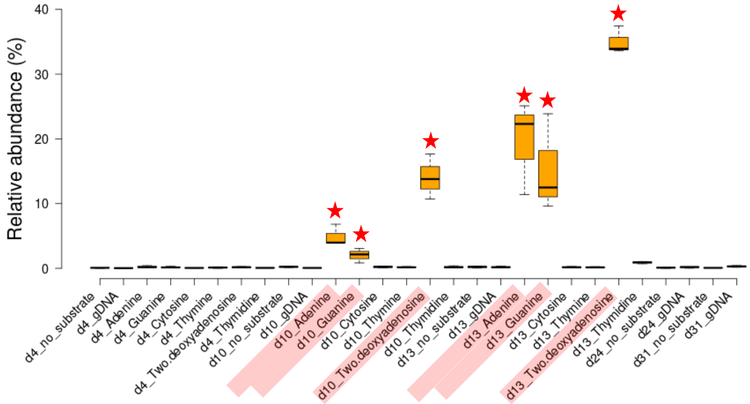


OTU 16:

OTU_16: Bacteria;Firmicutes;Clostridia;Clostridiales;Family_XII;Fusibacter;

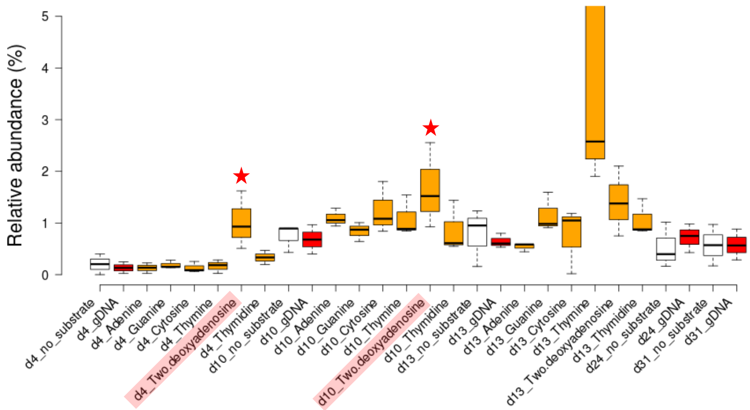


OTU 17:

OTU_17: Bacteria;Proteobacteria;Gammaproteobacteria;Alteromonadales;Shewanellaceae;Shewanella;

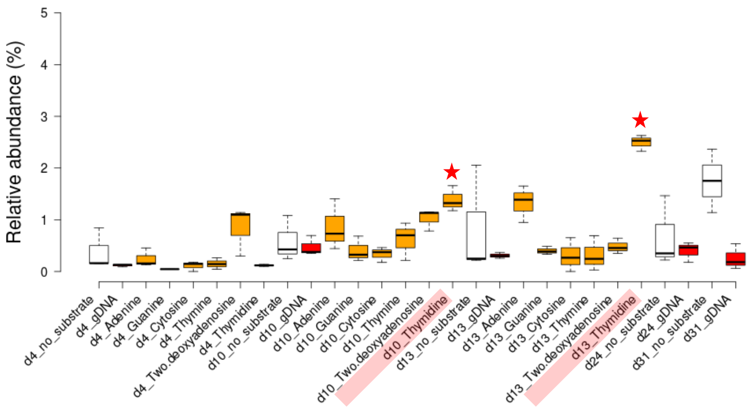

OTU 23:

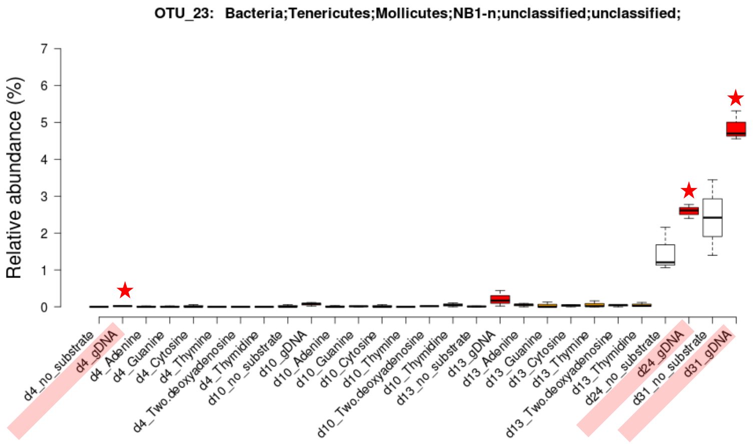


OTU 25:

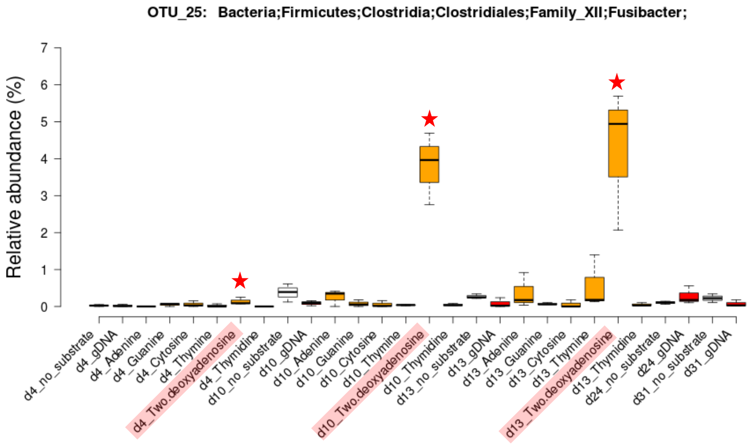


OTU 31:

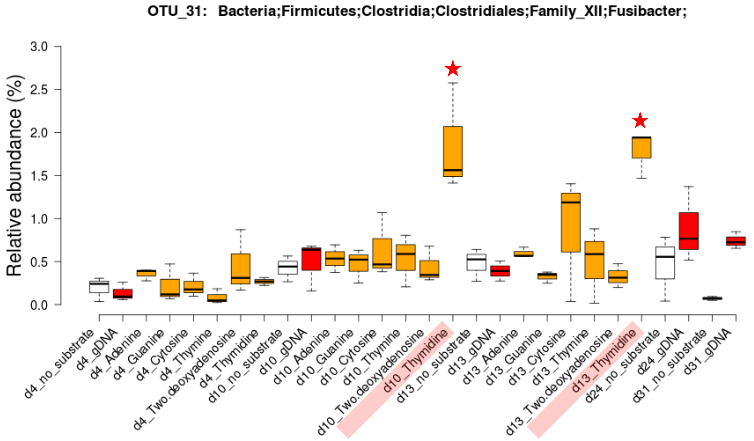


OTU 36:

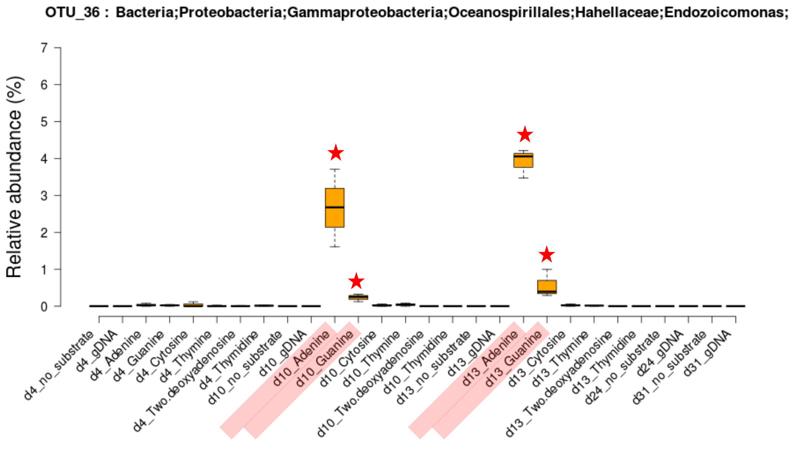

**OTU 48:**

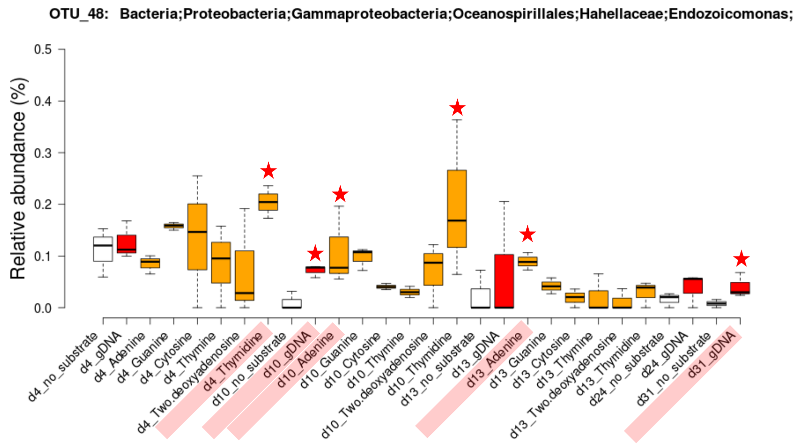


**OTU 55:**

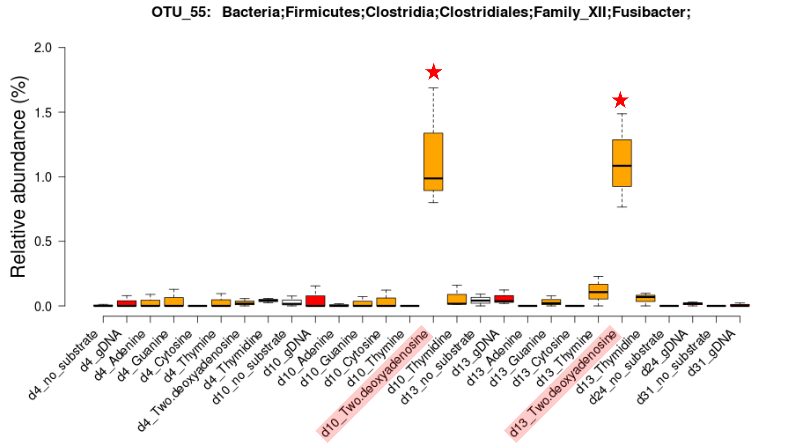


**OTU 60:**

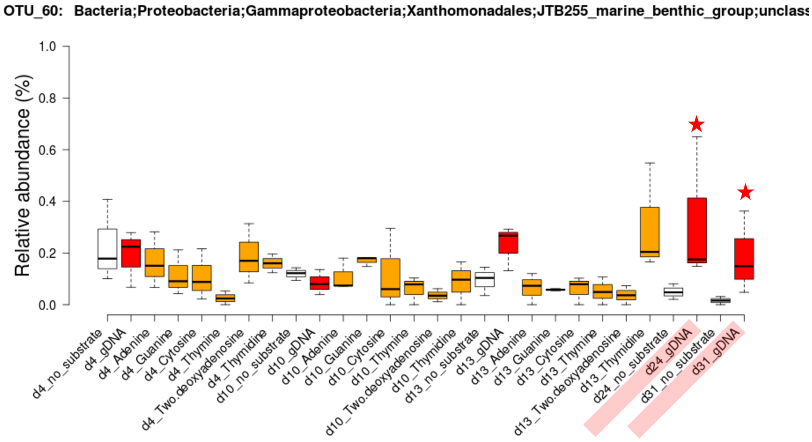


**OTU 70:**

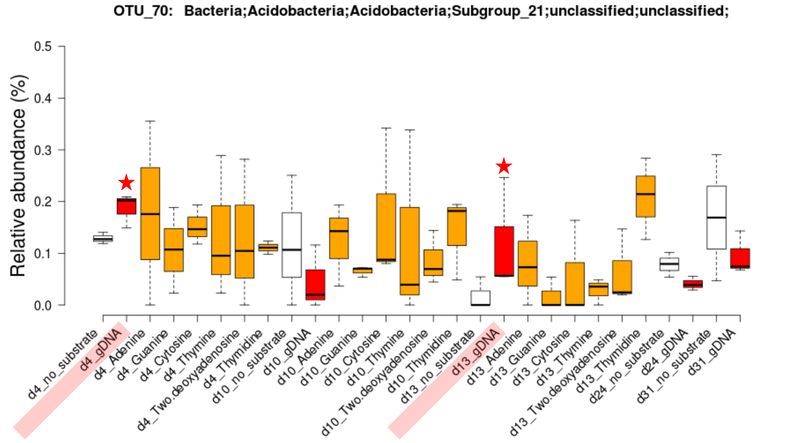

**OTU 86:**

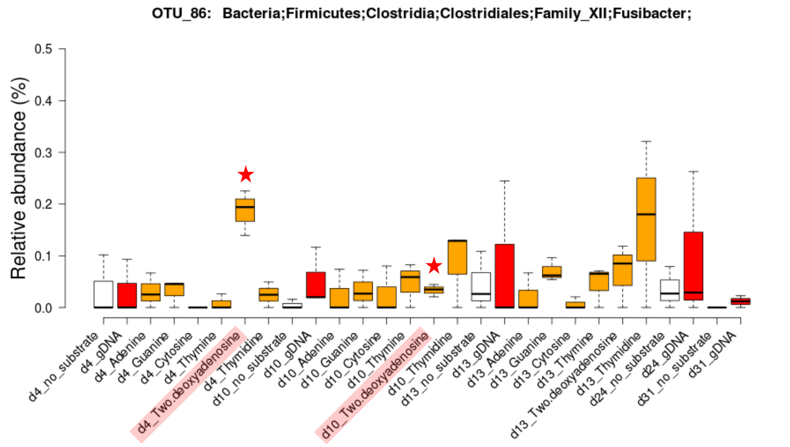


**OTU 140:**

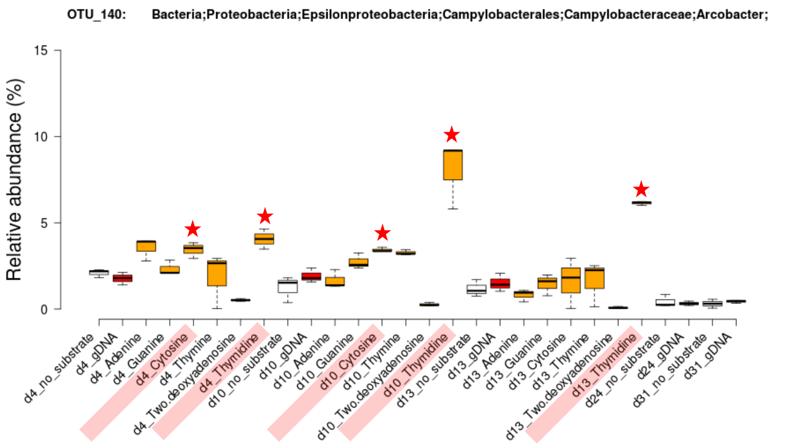


**OTU 144:**

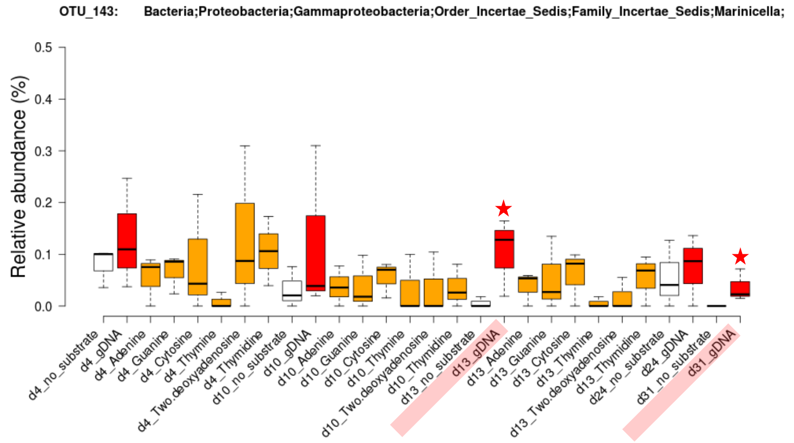


**OTU 167:**

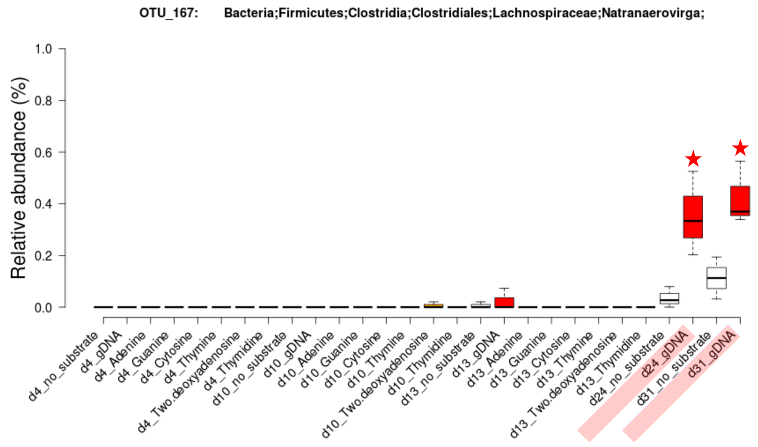

**OTU 204:**

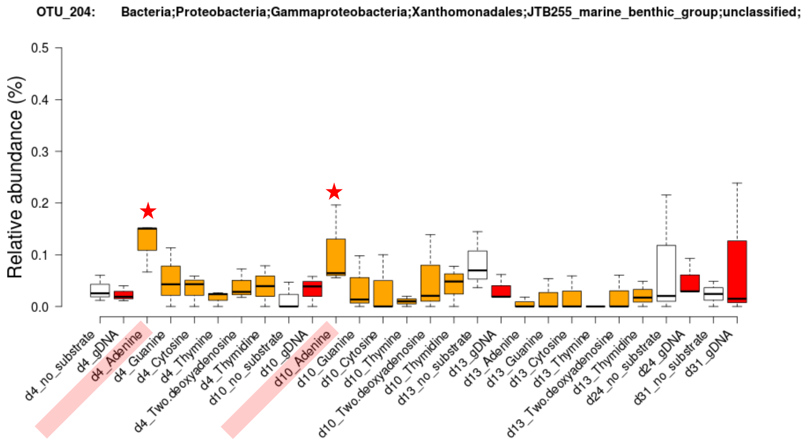


**OTU 219:**

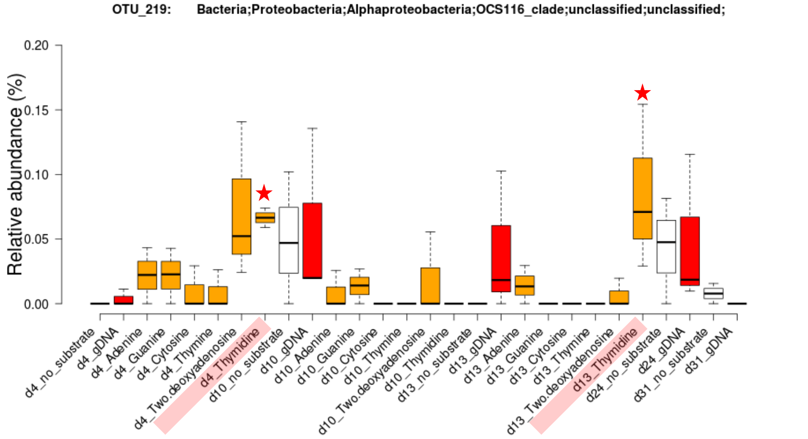


**OTU 224:**

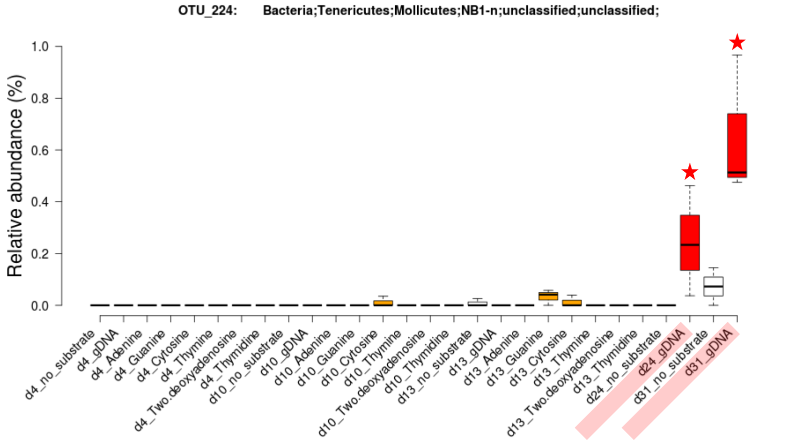


**OTU 255:**

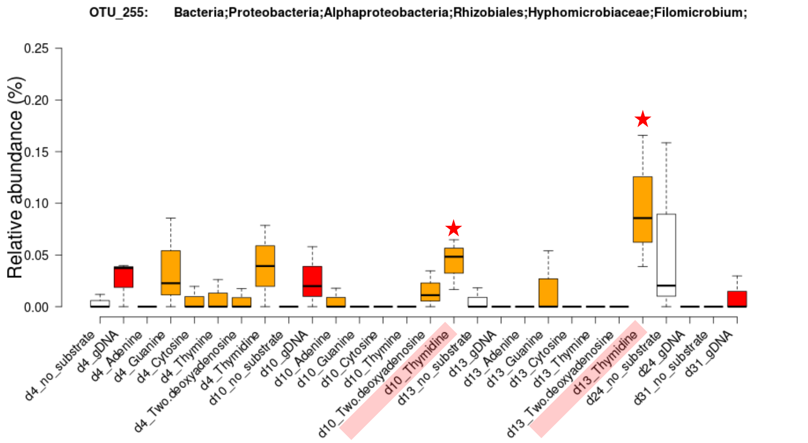

**OTU 310:**

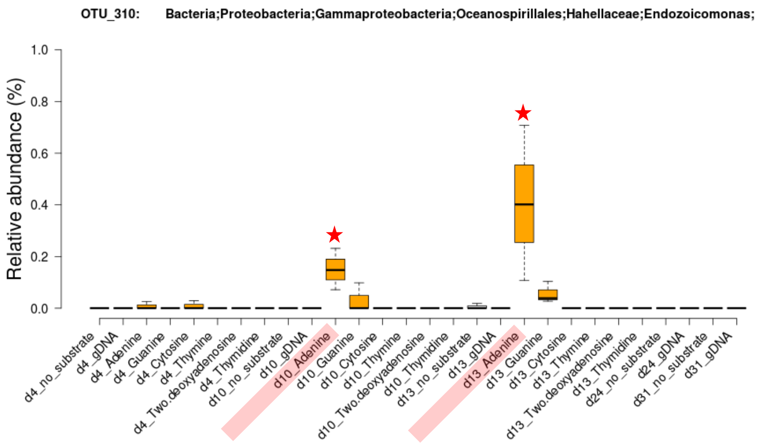


**OTU 515:**

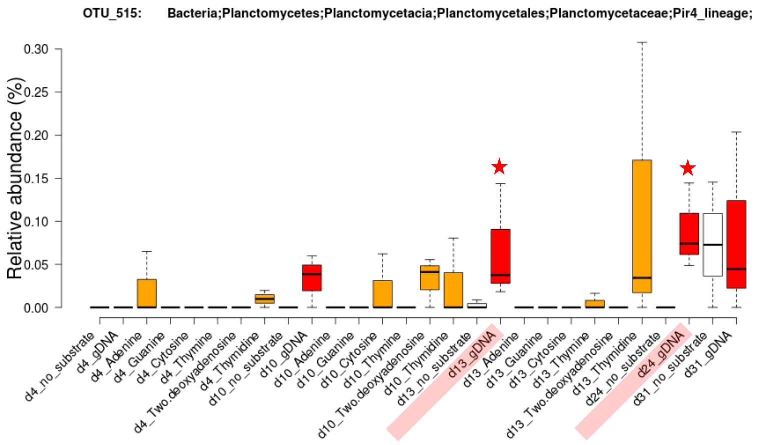


**OTU 810:**

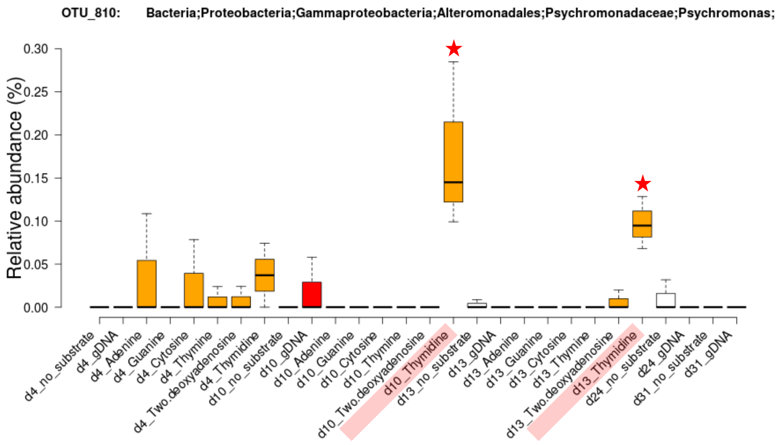


**OTU 1103:**

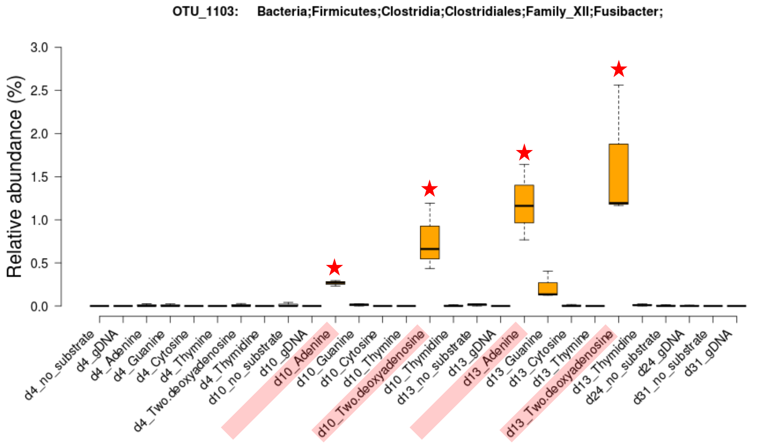

**OTU 1217:**

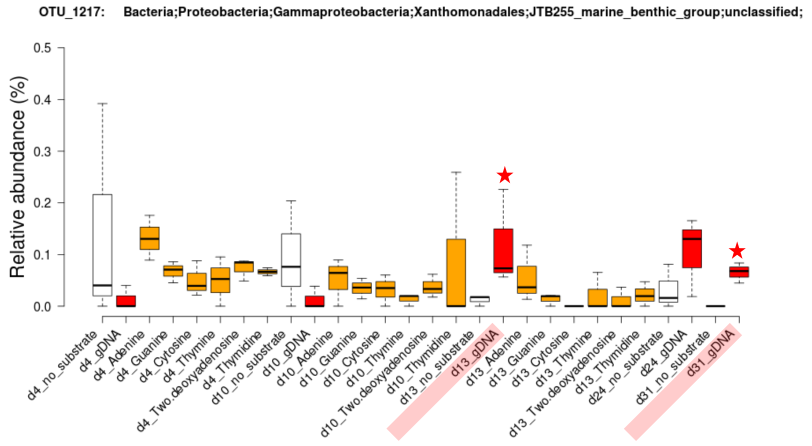


**OTU 1928:**

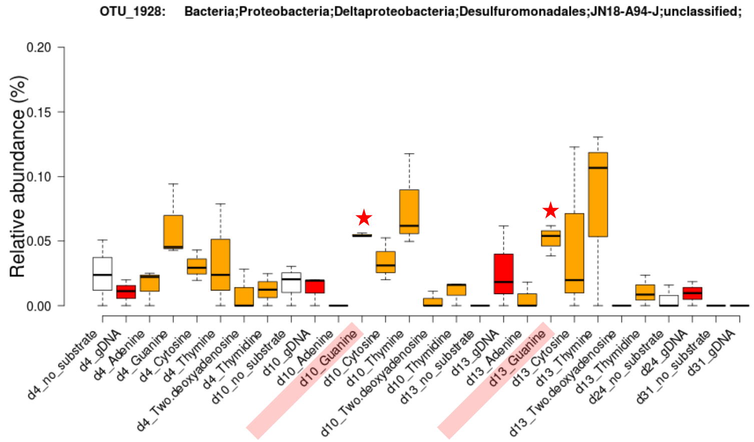


**OTU 2280:**

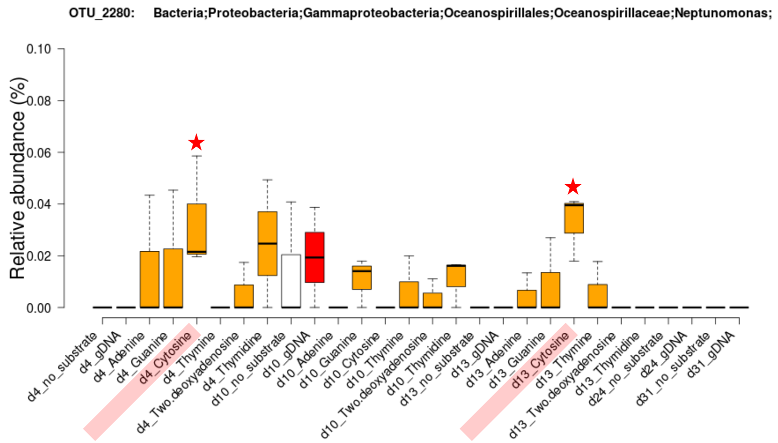


**OTU 4089:**

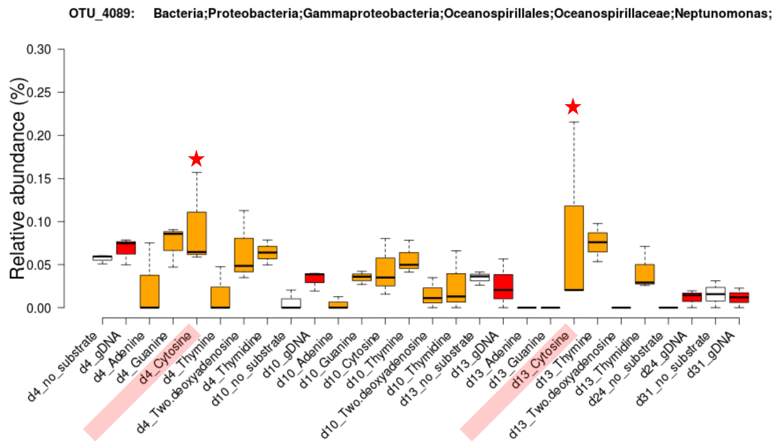

**OTU 4319:**

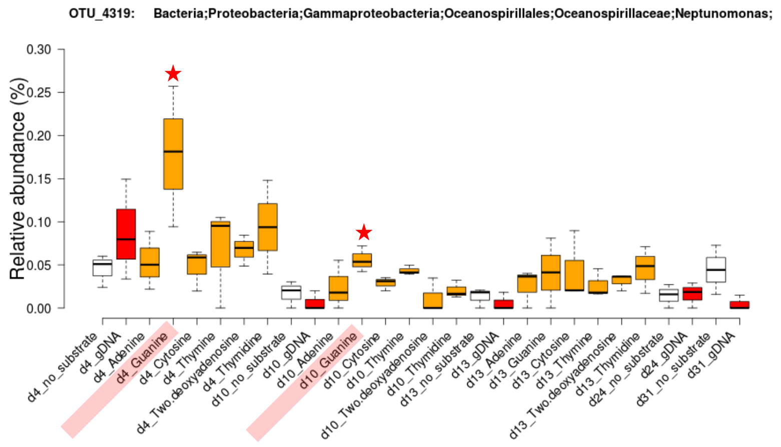


**OTU 4402:**

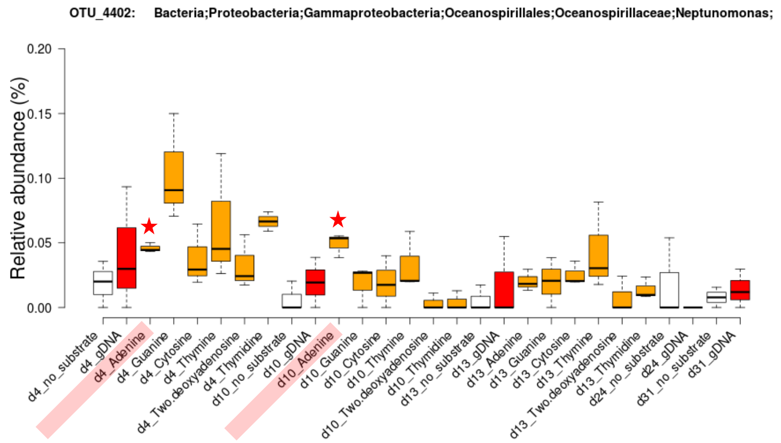


**OTU 4458:**

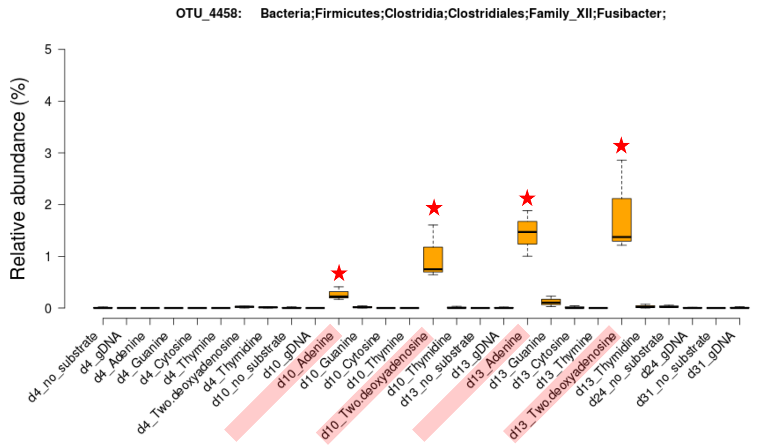


**OTU 6072:**

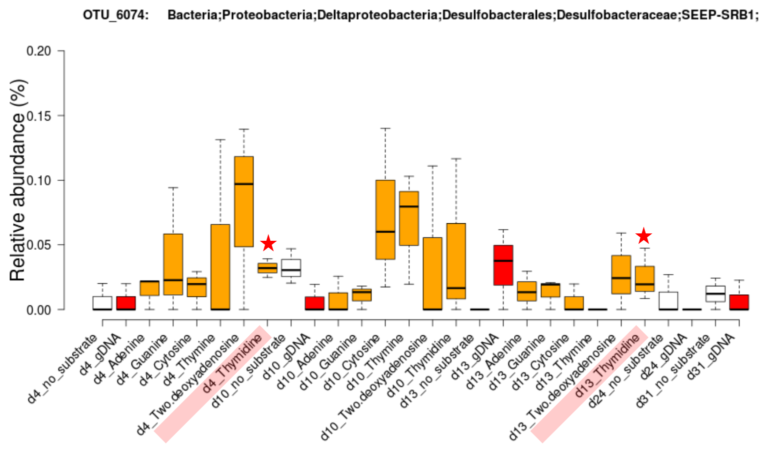

OTU 7362:

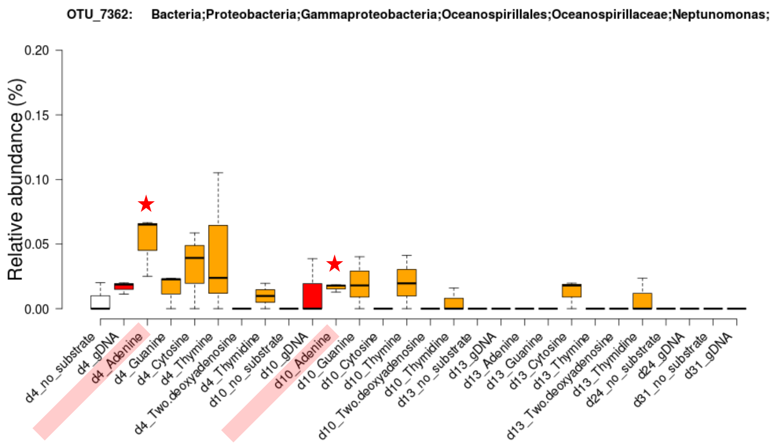


OTU 7702:

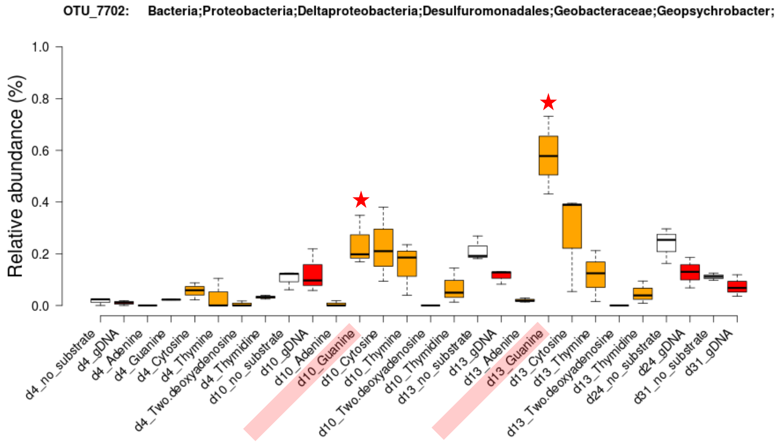


OTU 8132:

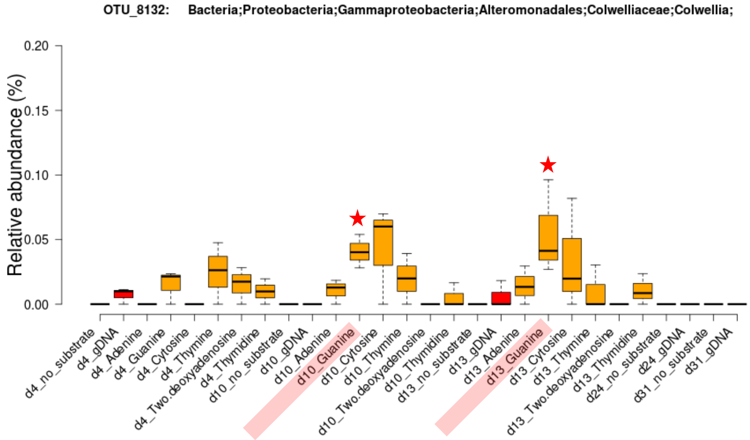
