## Supplementary material for "DNA-foraging bacteria in the seafloor": Supp. Fig. 9 - Xanthine dehydrogenase operon synteny

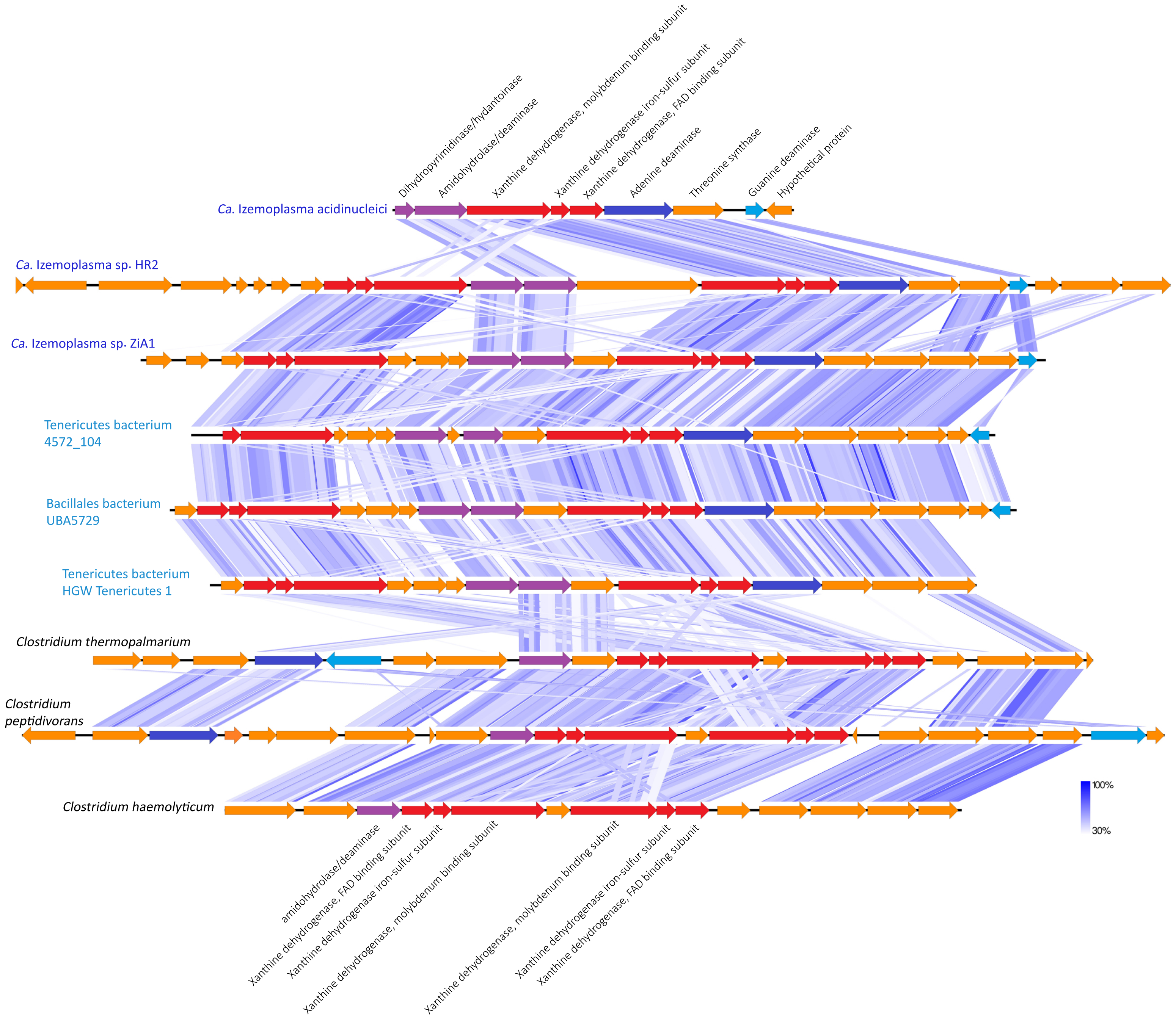


**Supp. Fig. 9.** Schematic of gene arrangments of 'xanthine-degradation loci' in *Ca.* Izemoplasmatales MAGs and selected *Clostridium* spp. genomes. Shaded blue lines show regions with high sequence similarity and synteny as determined by tBLASTx using EasyFig.
