## Supplementary material for "DNA-foraging bacteria in the seafloor": Supp. Fig. 10 - Halobacterium in SIP gradients

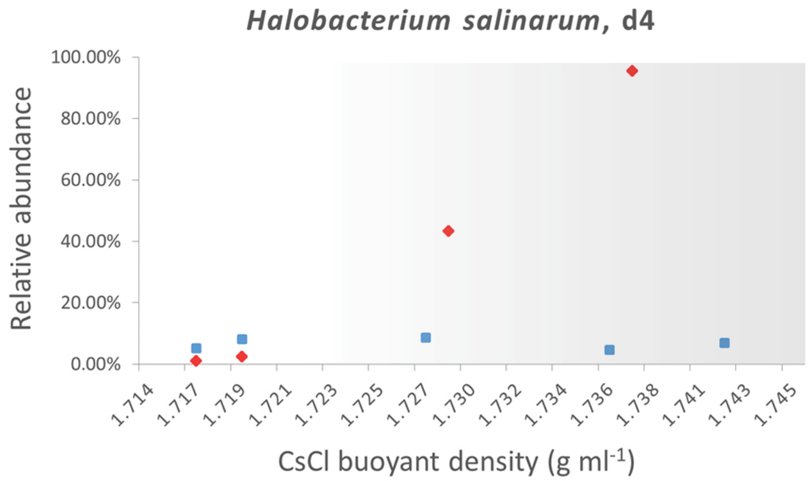


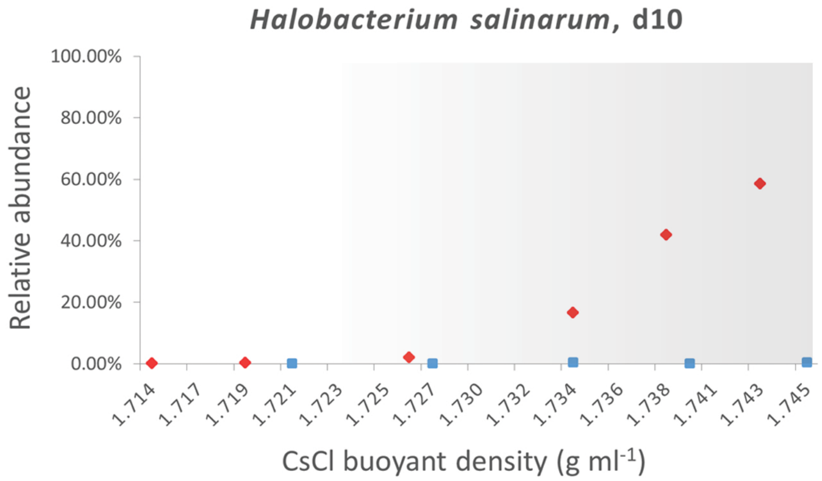


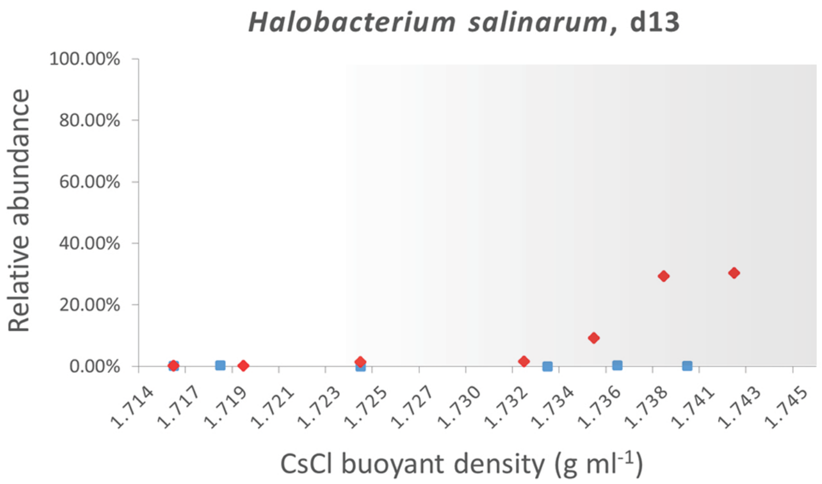


**Supp. Fig. 10.** Relative abundances of archaeal *Halobacterium salinarum* 16S rRNA genes across isopycicnic density gradient fractions, as determined by MiSeq amplicon sequencing. Relative abundances determined across $^{13}C$ and $^{12}C$ gradients are plotted in red diamonds and blue squares, respectively. The grey shaded areas represent the ‘heavy’ ends of the gradients.
