## Supplementary material for "DNA-foraging bacteria in the seafloor": Supp. Fig. 11 - BLASTP domain result

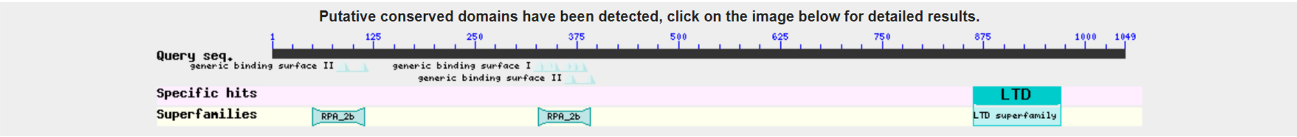


**Supp. Fig. 11**. Result of BLASTP analysis of peg.646 (Supp. Table 2) from *Candidatus* Izemoplasma acidinucleici against the Conserved Domain Database (CDD).
